# B-Cells Extracellular Vesicles Shape Melanoma Response to Immune Checkpoint Therapy

**DOI:** 10.1101/2024.12.12.628150

**Authors:** Ala’a Al Hrout, Agshin Balayev, Karla Cervantes-Gracia, Konstantinos Gkelis, Stephan Benke, Julia M. Matínez Gómez, Karina Silina, Mitchell P. Levesque, Richard Chahwan

## Abstract

The immune tumor microenvironment (TME) is increasingly recognized as a dynamic ecosystem where B cells play pivotal roles in modulating therapeutic responses, particularly in the context of immune checkpoint blockade (ICB) therapy. While B cells have traditionally been viewed as bystanders in tumor immunity, recent evidence suggests they may actively influence anti-tumor immunity, albeit with conflicting reports regarding their pro-tumor or anti-tumor roles. This study explores the crucial roles played by B cells and their secreted extracellular vesicles (EVs) in shaping melanoma responses to ICB therapy. We show a significant enrichment of B cells in ICB therapy responders compared to non-responders, pre-treatment, through retrospective analyses of melanoma patient tumors. Functional assays demonstrate that B cell depletion impairs T cell-mediated tumor cytotoxicity, underscoring the importance of B cells in anti-tumor responses. To investigate the clinical relevance, EVs were isolated from melanoma patient tumors, and fractioned into tumor and immune subpopulations. MiRNA profiling of CD19+ EVs identifies miR-99a-5p as a top candidate, among several others, upregulated in responders. Functional assays show that miR-99a-5p silencing in B cells diminishes T cell-mediated anti-tumor activity, suggesting its role in promoting B cell-mediated immune responses. Mechanistically, miR-99a-5p influences B cell maturation within the TME by mediating class-switch recombination. Our findings highlight the important role of B cells and their derived EVs in shaping the efficacy of melanoma immunotherapy, paving the way for novel therapeutic strategies targeting B cell-related pathways.

**Graphical abstract (created with Biorender):** 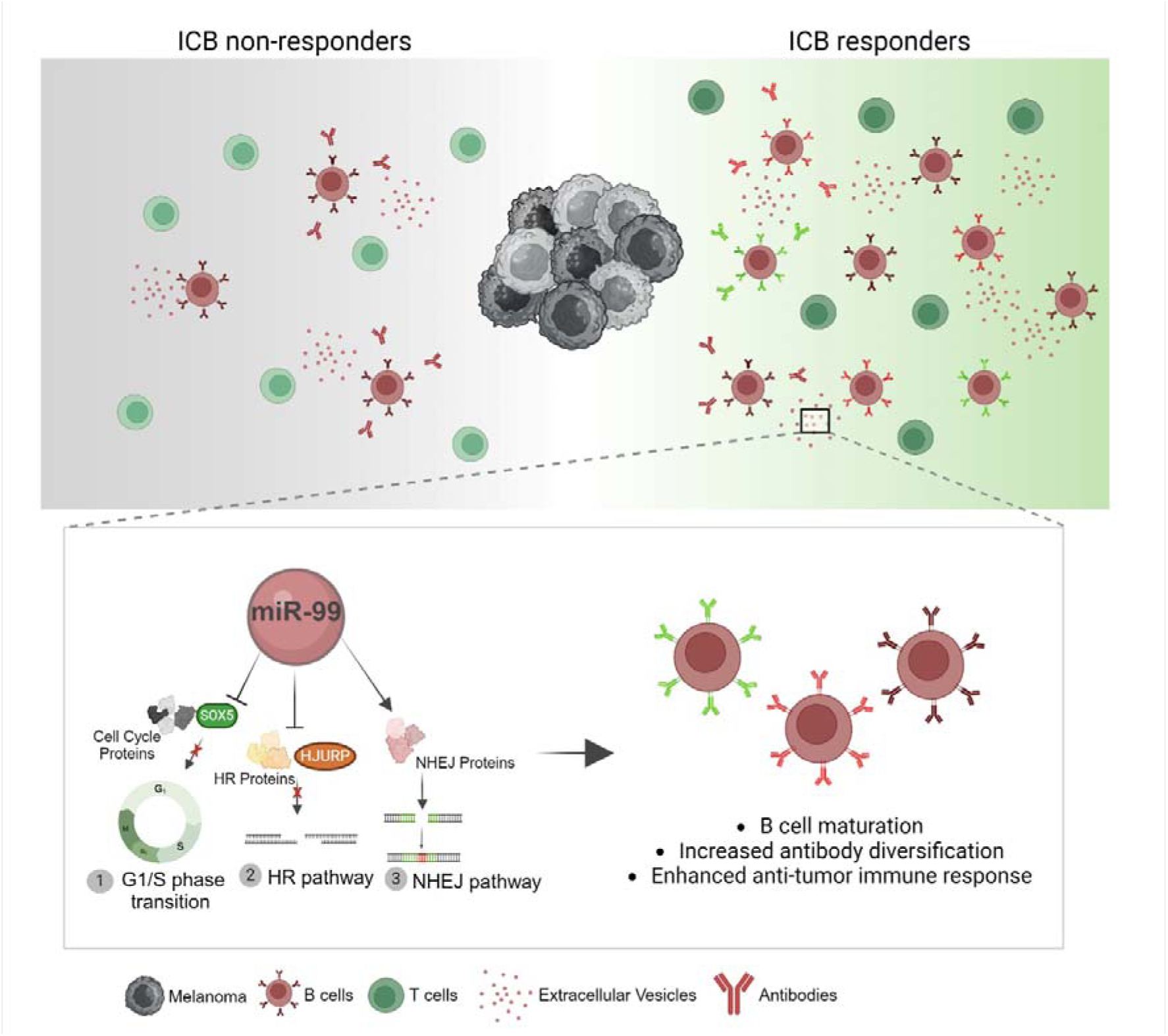

## INTRODUCTION

Immune cells, innate and adaptive, represent an essential component of the tumor microenvironment (TME), shaping cancer therapy outcomes. Their dichotomous properties hinge on the balance of tumor-promoting versus tumor-suppressing functions. T cells have long been considered paramount in context of immunotherapy and patient outcome due to their direct anti-tumor cytotoxicity. T cells represent the most common tumor infiltrating lymphocytes (TILs) and their enrichment in tumors has been correlated with favorable prognosis *(1–3)*. However, T cells also depend on B cells for efficient tumor-targeted cytotoxicity. B cells contribute to antitumor immunity through antibody production, cytokine secretion, antigen presentation. and co-stimulation of auxiliary immune cells *(4–6)*. Like T cells, tumor-infiltrating B cells (TIBs) could differentiate into several subtypes in response to cues from the TME. The understanding that B cells are a diverse population with a vast range of functions and properties can clarify their dual role in tumor immunity.

In melanoma, B cells can constitute up to half of the TILs population (reviewed in *(7)*). Studies have shown that they play an essential role in fostering an anti-tumor microenvironment and in activating T cells *(8)*. The contributions of B cells in immunotherapy highlight an unexpected role of this overlooked immune cell population in shaping the TME. Several studies reported a strong correlation between B cells enrichment and localization with immune-checkpoint blockade (ICB) therapy outcome in melanoma *(9–12)* The presence of B cells in melanoma biopsies prior to ICB treatment correlated with positive outcomes *(10, 11)*. And their colocalization with CD8^+^ T cells was linked to improved overall survival, independent of other clinical variables *(10)*. Tumor-associated B cells can localize within tertiary lymphoid structures that highly resemble secondary lymphoid organs and lymph nodes in their formation and organization *(7)*. Nonetheless, Tumor-associated B cells can also localize within the stroma or tumor margins, hinting that such B cell mediated effects could be achieved, in part, by proxy signaling. This could be mediated by classical B cell signaling such as antibody secretion and cytokines, but an additional enticing possibility is the involvement of extracellular vesicles (EVs).

EVs are emerging as complex signaling mediators within the TME. Tumor-derived EVs have been implicated in several stages of the tumorigenesis process, by regulating proliferation, angiogenesis, and immune-suppression *(13)*. EV mediated resistance has been described in melanoma through shuttling of tumor-derived PDL-1 on EVs *(14–17)*, suggesting a significant contribution of EVs in shaping immunotherapy outcome. Conversely, immune cells can regulate each other via EV-mediated signaling to promote immune-activation or immune-suppression based on TME cues. EV cargo contains proteins and small non-coding RNA (sncRNA) including miRNAs. Like gene expression, miRNAs can be differentially expressed between cells, hence, EVs-derived miRNAs also reflect that difference among different cell populations; even between EVs and their parent cells *(18)*. This suggests that EV cargo packaging and their transport is not a passive process, but an active and controlled one. As such, we postulate that EVs can carry signature miRNAs that can elucidate the contribution of certain cell populations in the context under investigation, for example, ICB therapy response. Despite the exciting studies reporting B cells role in promoting immunotherapy, most studies done are retrospective, relying on outcome post therapy. To gain more understanding, it is necessary to move from clinical data to mechanisms, which can ultimately translate into prognostic and therapeutic benefits. In this study, we explore the role of B cell-derived EVs in shaping the TME of ICB responders vs non-responders. We studied the *in situ* EVs profile from melanoma patients undergoing ICB therapy – from a cohort collected at baseline – through isolating tissue-derived EVs. EVs faithfully recapitulate the TME, where ICB therapy responders and non-responders can be divided into 2 cohorts based on their EVs surface markers and cargo, as if EVs had a “fingerprint” that convey their origin and microenvironmental context. Nonetheless, bulk EVs sequencing can be misleading, or at best, uninformative when looking at subtle changes in specific subpopulations within the whole EV population. Therefore, melanoma and B cell-derived EVs were individually isolated from the heterogenous mixture of bulk EVs and their miRNA cargo was isolated and sequenced. Strikingly, the EVs landscape reflected what we and others have observed on the cell level, of enrichment of immune cell infiltration in responders over non-responders. B cell-derived EVs had far more unique miRNAs that are not shared with melanoma-EVs; highlighting that B cells exhibit significant change in context of ICB therapy. miR-99a-5p, a well-established tumor suppressor, was identified in B cell-derived EVs from responders. In downstream functional assays we were able to show that miR-99a-5p exhibit a B cell-dependent phenotype, where it regulates the cell cycle and DNA damage repair pathways, resulting in enhanced CSR. We propose that B cells in ICB therapy responders exhibit higher CSR and enhanced anti-tumor response via EV-based signaling, in part, via miR-99a-5p. These findings collectively suggest that B cells and their derived EVs are remarkable contributors to shaping therapeutic outcome in context of ICB therapy in melanoma.

## RESULTS

### B cells are enriched in tumors of ICB therapy responders vs. non-responders pre-treatment

Our meta-analysis of large scale RNAseq data from 3 independent melanoma studies *(19– 21)*(; total n=153) at baseline, prior to undergoing ICB therapy, provided insights into the role of B cells in shaping the melanoma response to immunotherapy by comparing responder (n=81) vs non-responder (n=72) cohorts (Fig. 1A). The estimation of immune infiltration in ICB responders over non-responders revealed a significant enrichment of B cells (Fig. 1B). This unexpected finding was further corroborated by analyzing differentially expressed genes (DEGs) between responders and non-responders. The top 50 significant DEGs are shown in Fig. 1C. Significantly upregulated DEGs in ICB therapy responders predominantly included genes specifically expressed by B cells and/or influencing B cells development and homing, including CD19, AICDA, PAX5, and BLK (Fig. 1C, bold). Validation in a fourth independent dataset of SingleCell RNAseq data of melanoma tumors *(22)* visualized on <u>Single Cell portal</u> (Broad Institute) corroborated these findings; expression of significant upregulated DEGs from our meta-analysis was enriched in the B cells cluster, rather than the melanoma cluster (Fig. 1D-E). Furthermore, protein-protein network analysis of significant upregulated genes (encircled in yellow) and their immediate protein neighbors highlighted proteins that shape B cells responses as a major contributor to the network (Fig. 1F). These proteins (in red) play a role in B cell activation, differentiation, somatic recombination of immune receptors (immunoglobulin superfamily), and B cell receptor (BCR) signaling (Table S1). To understand the function of the upregulated genes in responders, we carried out gene ontology (GO) analysis of biological process term enrichment. As predicted, and in concordance with our earlier results, the enriched terms generally included immune response regulation; in addition to somatic recombination of immune receptors and B cell activation (Fig. 1G).

**Figure 1.**
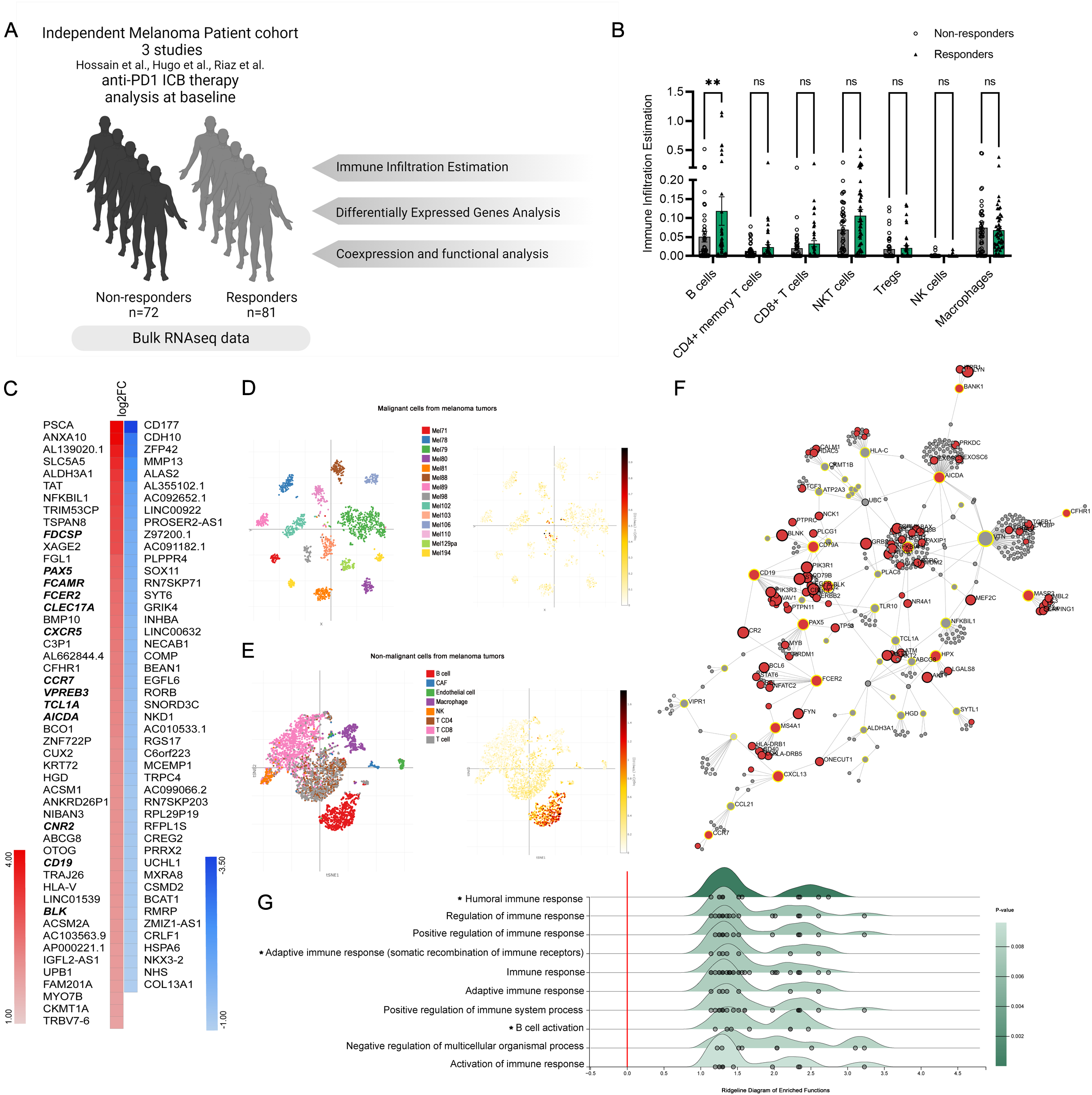
B cells are enriched in the tumors of ICB therapy responders in melanoma patient’s cohort. (A) Schematic depiction of datasets design for meta-analysis. (B) Immune infiltration estimation based on bulk RNAseq data. Estimation was done using TIMER2.0. *P* values were determined by two-way ANOVA with Sidak multiple comparison test (p <0.004). (C) Expression heatmap of the top 50 significant DEGs in responders in comparison to non-responders. (D) tSNE plot of scRNAseq of malignant cells from melanoma tumors and expression location of the top 100 significant upregulated genes *(22)* visualized on <u>Single</u> <u>Cell portal</u> (Broad Institute). (E) tSNE plot of scRNAseq of non-malignant cells from melanoma tumors and expression location of the top 100 significant upregulated genes visualized on <u>Single Cell portal</u> (Broad Institute). (F) protein-protein interaction network of significant upregulated genes in responders over non-responders (encircled in yellow), and their immediate network. Proteins in red are involved in BCR signaling pathway (full list in table S1). Network is visualized using NetworkAnalyst *(70)*. (G) Ridgeline diagram of enriched biological process of the significant upregulated genes. X-axis shows expression fold change of responders over non-responders. Ridgeline analysis was done with ExpressAnalyst *(68)*.

### T cell anti-tumor cytotoxic activity is impaired by B cell depletion

To address our initial observation that B cells might play an important role in T cell mediated tumor killing in melanoma, we developed a co-culture tumor-killing assay. In our experimental setup (Fig. 2A), we first label melanoma cells with cell trace (CT) to distinguish them from the remainder of the PBMC co-culture. Activation of PBMCs with Staphylococcal Enterotoxin B (SEB) crosslinks TCR with MHC-II on antigen presenting cells and bypasses the need for specific TCR-MHC-peptide complex interaction to elicit a strong immune response *(23–25)*. We then compare cell death of melanoma cells (positive for Annexin V/ 7AAD) when co-cultured with complete or B cell-depleted PBMCs (Fig. 2B, top panel). Interestingly, B cell depletion resulted in a significant decrease in melanoma’s cell death in comparison to complete PBMCs culture (Fig. 2B, bottom panel), without affecting CD3^+^ T cell numbers (Fig. S1A). This suggests that T cells anti-tumor cytotoxic activity might be impaired by B cell depletion in melanoma samples.

**Figure 2.**
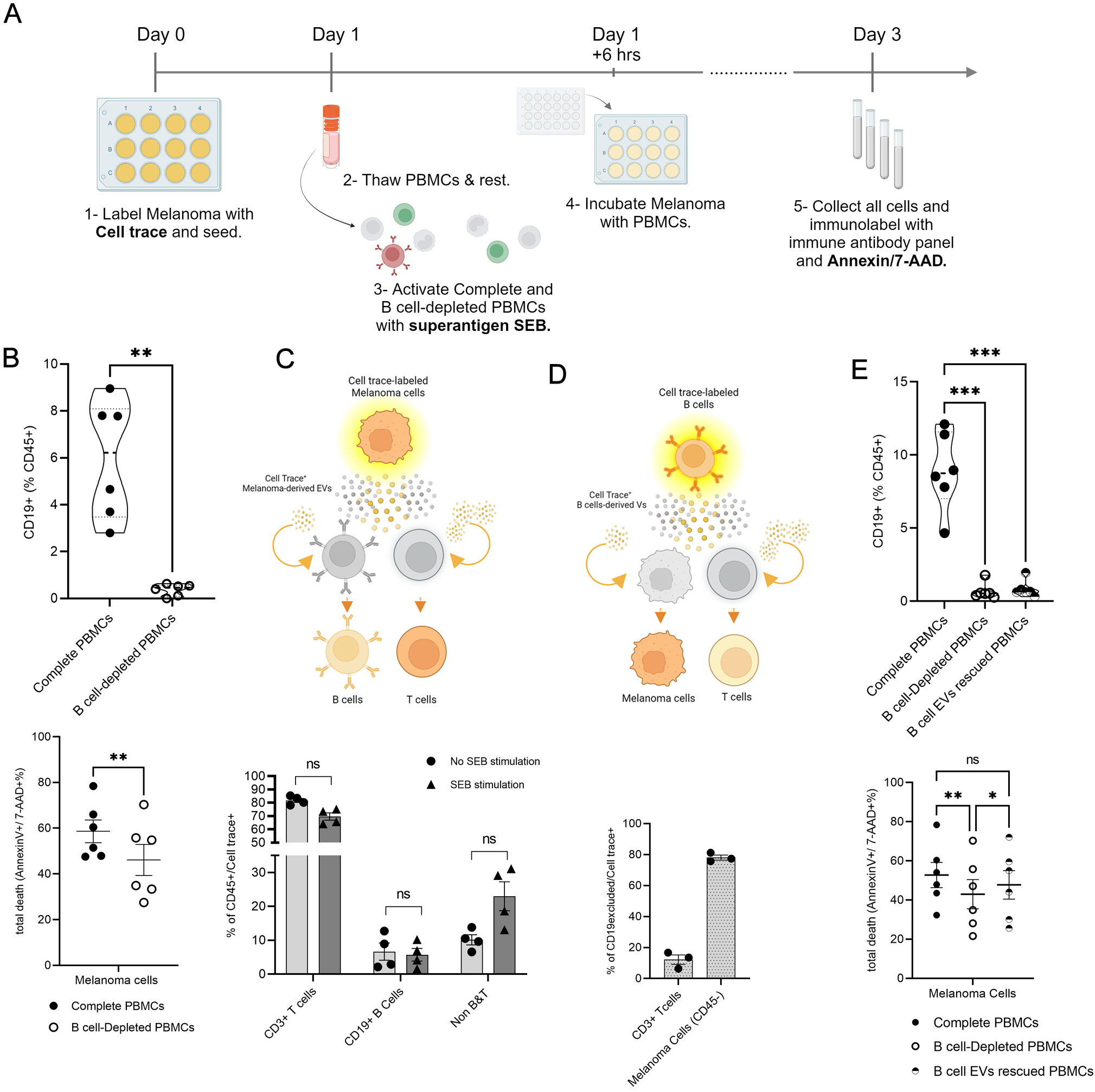
Cell-cell signaling is mediated through EVs signaling. (A) Schematic depiction of tumor-killing assay setup. (B) CD19^+^ cells in complete and B cell-depleted cultures. Presented as percentage of CD45^+^ population. *P* values were determined by two-tailed paired t-test (**p<0.005). Tumor-killing assay in complete and B cell-depleted cultures, presented as total death of melanoma cells that are single and double positive to Annexin-V-FITC and 7-AAD. *P* values were determined by paired t-test (**p<0.005). (C) EVs uptake from cell-trace-stained melanoma cells. CD3^+^, CD19^+^, CD3^-^CD19^-^ populations out of CD45^+^/Cell trace^+^ with and without SEB-stimulation, presented as percentile. *P* values were determined by Sidak multiple comparison t-test. (D) EVs uptake from cell-trace-stained B cells. Melanoma population and CD3^+^ population out of CD19excluded^/^/Cell trace^+^ without SEB-stimulation, presented as percentile. (E) CD19^+^ cells in complete, B cell-depleted, B cell EVs-rescued cultures. Presented as percentage of CD45^+^ population. *P* values were determined by one-way ANOVA (***p<0.001). Tumor-killing assay in complete, B cell-depleted, B cell EVs-rescued cultures, presented as total death of melanoma cells that are single and double positive to Annexin-V-FITC and 7-AAD. *P* values were determined by one-way ANOVA (*p<0.05, **p<0.01).

### TME cell-cell signaling can be mediated by EVs

In our tumor killing assay setting, we observed a CD45^+^ population that is, unexpectedly, also positive for CT used to exclusively stain melanoma cells (Fig. S1B). Since CT is quickly inactivated by cell culture media it could not have escaped from stained melanoma cells to immune cells. Yet some of the expected CD45+/CT-population have uptaken CT stain from melanoma and became CD45^+^/CT^+^. Much of this double positive population were CD3^+^ T cells, but CD19^+^ B cells were also stained (Fig. 2C, S1C). Similarly, we isolated B cells from PBMCs and tagged them with CT to be co-cultured with the rest of PBMCs and melanoma cells (Fig. 2D, S1D). The majority of non-B cells positive for CT were melanoma cells, suggesting an active and deliberate, rather than passive, CT uptake (Fig. 2D). This is in line with the ability of EVs to transfer cargo across cells especially in the TME *(26)*. To address this, we validated that secreted EVs from CT-tagged cells are themselves CT^+^ (Fig. S1E). To further confirm the possible role of B cell derived EVs in T cell mediated anti-tumor activity in the tumor killing assay, we depleted PBMCs from B cells and then rescued the culture with B-cell derived EVs (Fig. 2E). Rescuing PBMCs with B cell-derived EVs was sufficient to reinstate tumor killing ability even in the absence of B cells (Fig. 2E).

### In-situ EV analyses highlight differences between ICB responders vs non-responders

To better replicate in vivo and in situ EV-based signaling within the TME we explored the isolation of EVs from patient-derived tumor tissue; we had previously shown that melanoma biopsies possess abundant tumor and immune EV particles *(27)*. With this added approach we would be able to further fractionate and analyze EVs derived from the different TME cell populations resident at the tumor site at the time of biopsy sampling. Using flow cytometry to quantify cells directly from our melanoma biopsy cohort (n=10) we observed a significant enrichment of B cells in melanoma biopsies from ICB therapy responders vs non-responders (Fig. 3A). This result supported our metanalysis of the 3 independent melanoma datasets. Tissue-derived EVs were then isolated, processed, and imaged using transmission election microscopy (TEM) to verify their size and morphology (Fig. 3B-C), as elaborated in our prior work *(27)*. The isolated particles displayed expected EV characteristics including: i) size, ranging between 50-180 nm and ii) a round concaved bi-layer morphology (Fig. 3B-C). After we established the successful isolation of EVs, we carried out size estimation of the particles using nano-flow cytometry *(27–29)*. NanoFCM sized the isolated EVs accurately, binning them into sub-populations of different sizes, reflecting the size range observed by TEM (Fig. 3D-E). The quality of our EV prep was further verified using nano-flow cytometry and quality controls to exclude background noise and non-specific staining (Fig. S2A). Our EV isolation displayed high purity, averaging between 60% and 90% as measured by Cell Trace, excluding non-biological debris (Fig. 3F). Staining for CD63 also showed high percentage, further confirming EV isolation efficacy (Fig. S2B).

**Figure 3.**
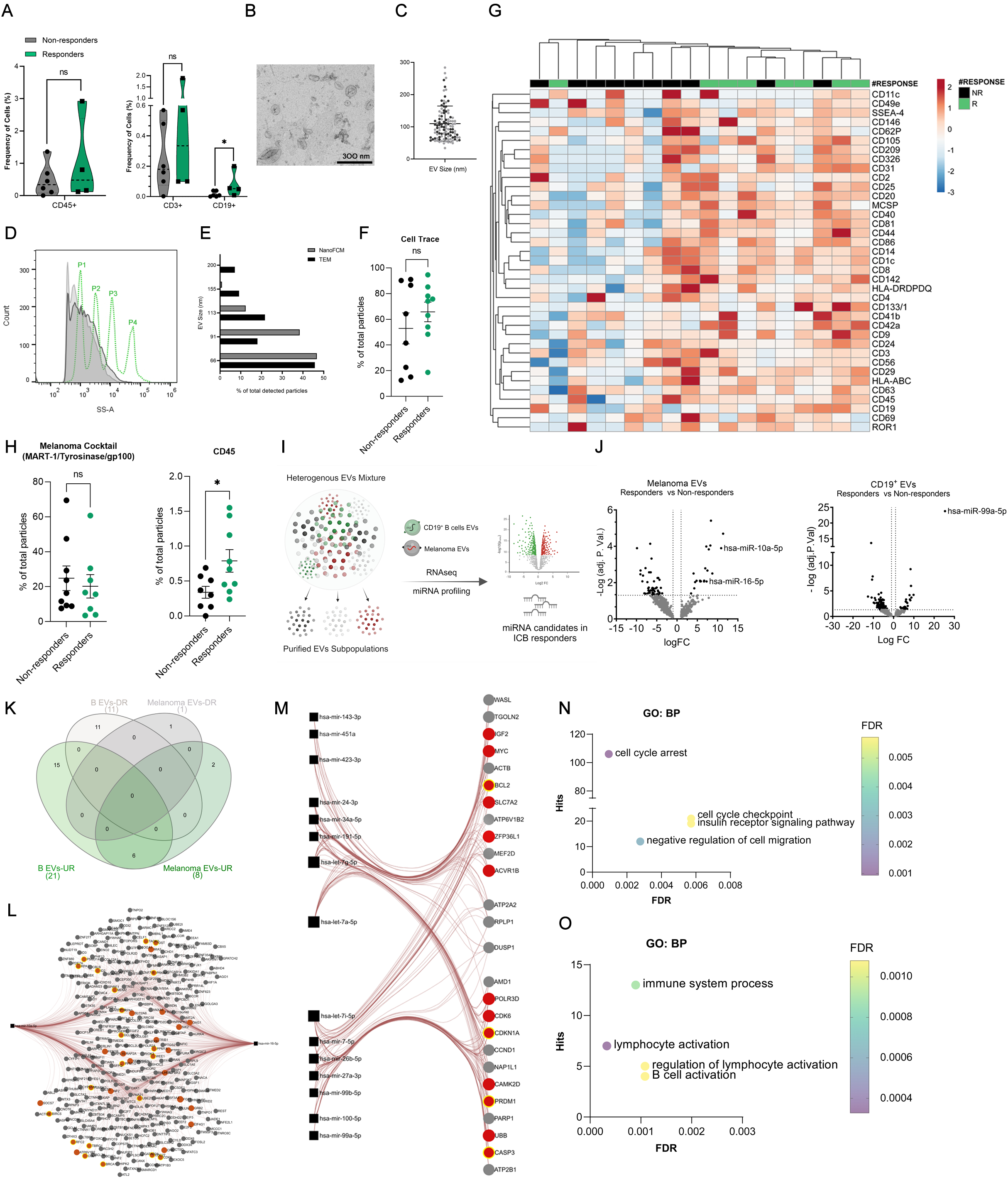
In-situ EVs analysis highlights differences between responders and non-responders to ICB therapy. (A) CD45^+^, CD3^+^, CD19^+^ populations in responders and non-responders processed tissue, presented as frequency of total cells. P values were determined by two-sided Mann–Whitney test U-test (*p<0.05). (B) TEM representative imgae of isolated EVs from patient tumor tissue biopsies (C) size distribution of detected particle visuzalized by TEM images. Analysis was done using MAPS software. (D) Histogram of size distribution of isolated EVs (light and dark gray, n=2) in comparison to silica sizing nanospheres (dotted green line). Samples were aquired using nano analyzer NanoFCM. (E) Comparison of EVs sizes obtained from TEM and nano-analyzer. (F) cell trace positive EVs, presented as percentage of detected particles in responders (green, n=9) and non-responders (black, n=8). Each point is an independent patient sample. (G) Heatmap of expression of denoted surface markers of tissue-derived EVs from responders (n=8) and non-responders (n=10). Origninal values are ln (x+1) transformed. Rows are centered, unit variance scaling is applied to rows. Rows are clustered using Euclidean dsitance and average linkage. Columns are clustered using maximun distance and average linkage. Created with ClustVis *(71)*. (H) Melanoma cocktail^+^ (MART-1, Tyrosinase, and gp100) and CD45^+^ EVs, presented as percentage of detected particles in responders (green, n=9) and non-responders (black, n=6). P values were determined by Welch’s t-test (*p<0.05). (I) schematic depiction of EVs fractionation setup. Created with Biorender. (J) volcano plots of DE miRNAs in responders (n=8) over non-responders (n=6) from melanoma and CD19^+^ derived EVs. (K) Venn diagram of significant upregulated and downregulated miRNAs in responders over non-responders from melanoma and CD19^+^ derived EVs. Created with InteractiVenn *(72)*. (L-M) Melanoma-derived EVs (L) and CD19^+^ EVs (M) unique upregulated miRNAs in responders and their predicted targets. Analysis was done with miRnet 2.0 *(67)*. (N-O) Gene enrichment analysis of selected biological processes associated with predicted gene targets of melanoma-derived EVs (N) and CD19^+^ EVs (O) unique upregulated miRNAs in responders.

Multiplex EV surface protein detection gave us a comprehensive overview of tissue-derived EVs in responders and non-responders, including immune cells-derived EVs. Overall, tissue-derived EVs from responders had a higher expression of immune markers (Fig. 3G). Interestingly, hierarchical-clustering analysis of the heatmap showed a segregation between responders and non-responders, highlighting that tissue-derived EVs recapitulated what we have observed thus far in the cellular TME of immune populations enrichment in responders over non-responders; underscoring the reliability of tissue-derived EVs in representing the in-situ TME. This observation was further corroborated with the analysis performed using nano-flow cytometry detection of melanoma and immune derived EVs on the single particle level. Immune cells-derived EVs (CD45^+^) were significantly higher in responders over non-responders (Fig. 3H), whereas melanoma EVs did not show a significant difference between the two cohorts, suggesting a substantial change occurs on immune cell-derived EVs rather than melanoma-EVs in the context of ICB therapy.

### In situ B cell-derived EVs are a differentiating factor in melanoma ICB response

Using the results we have achieved thus far, we aimed at untangling the complex network of EVs by fractionating the heterogeneous mixture of tissue-derived EVs into subpopulations based on their surface markers (Fig. 3I). The miRNA cargo of the subpopulations of interest (i.e. B cell- and melanoma-derived EVs) was profiled. We compared each EV subpopulation between responders and non-responders (e.g. CD19^+^ responders vs CD19^+^ non-responders). Based on differential expression analysis, miRNAs that met the cutoff threshold (Log_2_FC [≥1, FDR<0.05) were defined as significant differentially expressed (DE) miRNAs (Fig. 3J). When comparing ICB responders vs non-responders, 21 miRNAs were upregulated in CD19^+^ EVs, and 8 miRNAs in melanoma EVs. This might suggest that changes in miRNA cargo in response to ICB therapy are more pronounced in B cells-derived EVs than in melanoma cells. Therefore, we postulate that DE miRNAs that are unique to B cell-derived EVs could shed more light on the mechanistic role of B cells in shaping the response to ICB. As a result, DE miRNAs in responders vs non-responders of B cell-derived and melanoma-derived EVs were plotted in a Venn diagram (Fig. 3K).

We explored the predicted gene targets of the unique miRNAs identified to be upregulated in melanoma-derived EVs (Fig. 3L) and B cell-derived EVs in responders (Fig. 3M). Melanoma-derived EVs had only 2 unique miRNAs that are not shared with B-cell derived EVs. Predicted gene targets of these unique miRNAs were enriched in cell-cycle arrest and cell-cycle checkpoint pathways (Fig. 3N). The predicted targets of B cell-derived EVs miRNA were enriched in immune system process pathway, lymphocyte activation pathway, and B cell activation pathways (Fig. 3O). Given the scope of this study, we decided to focus on B cell-derived EVs miRNA for our mechanistic studies.

### miR-99a-5p of B cell EV promotes melanoma ICB response

miR-99a-5p was the top hit among the 15 unique miRNAs identified in B cell-derived EVs in responders. miR-99a has been established as a master tumor suppressor across many cancers *(30–33)*. However, the expressing cell and the mechanism by which miR-99a-5p exerts its tumor-suppressing function remains unknown. Herein, we propose that miR-99a-5p could play a critical role in the anti-tumor response B cells play in context of melanoma ICB therapy.

To that end, we treated PBMCs with miR-99a-5p silencer (antagomir-99a) or with negative control. We compared complete PBMCs and B cell-depleted PBMCs killing ability with silencing miR-99a-5p (Fig. 4A). Remarkably, silencing miR-99a-5p significantly impaired tumor-killing ability, and only in the presence of B cells where the strong phenotype was lost upon B cells depletion (Fig. 4A). Silencing miR-99a-5p significantly reduced the percentage of CD19^+^ B cells, but didn’t affect the percentage of CD3^+^ T cells (Fig. 4B-C).

**Figure 4.**
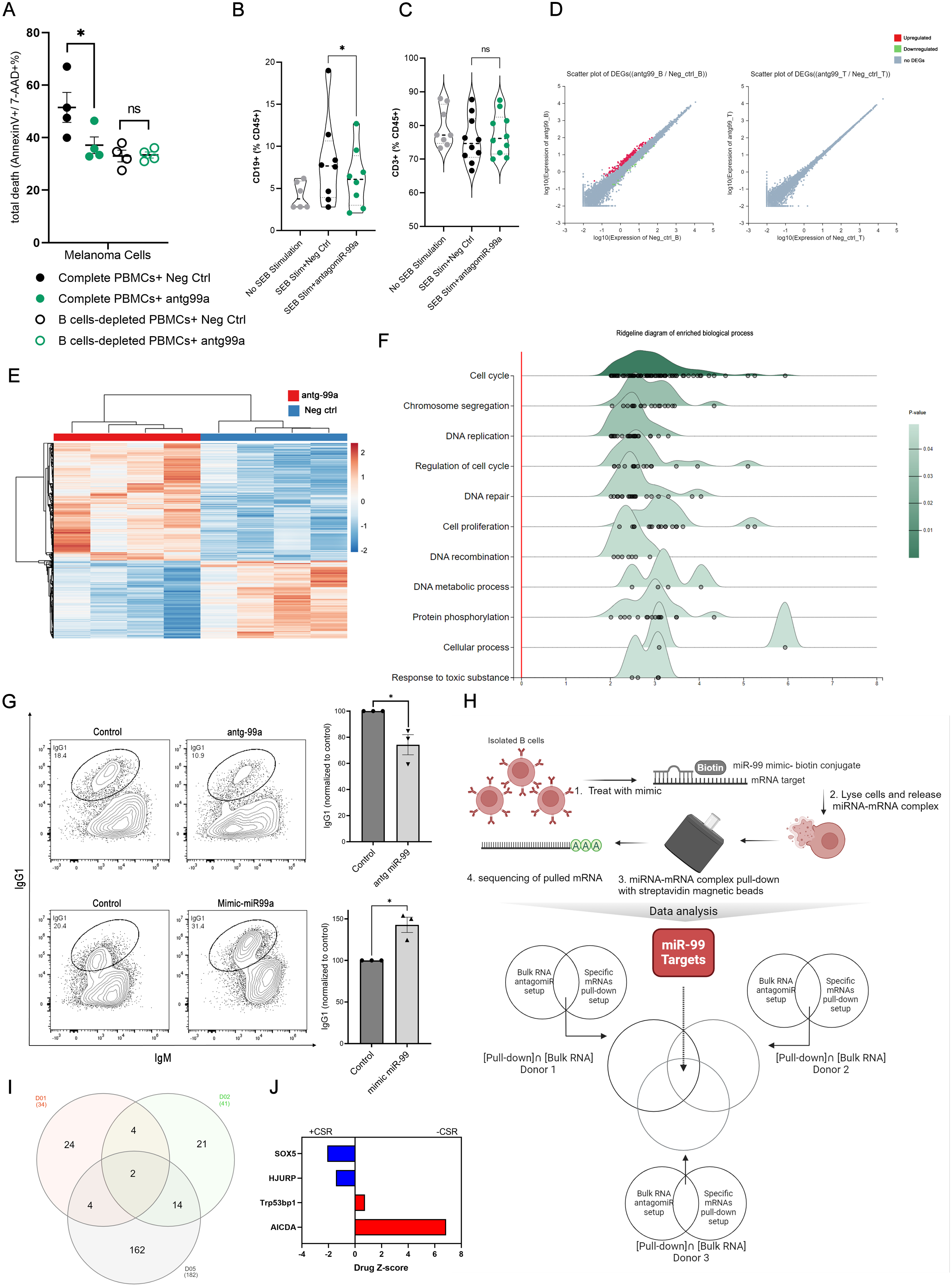
miR-99a-5p B cells mediated phenotype is through regulation of the cell cycle and CSR. (A) tumor-killing assay in complete and B cell-depleted cultures treated with antagomiR-99a-5p or negative control antagomiR, presented as total death of melanoma cells that are single and double positive to Annexin-V-FITC and 7-AAD. *P* values were determined by two-tailed paired t-test (*p<0.05). (B-C) CD19+ B cells (B) and CD3+ T cells (C) with or without SEB stimulation and treated with antagomiR-99a-5p or negative control antagomiR. Presented as percentage of CD45+ population. *P* value was determined by two-tailed paired t-test (*p<0.05). (D) Scatter plot of DEGs in isolated B cells and T cells treated with antagomiR-99a-5p or negative control antagomiR (n=4). (E) hierarchical heatmap clustering of transcriptome profiles of isolated B cells treated with antagomiR-99a-5p or negative control antagomiR (n=4). Rows are centered, unit variance scaling is applied to rows. columns are clustered using correclation dsitance and average linkage. Created with ClustVis *(71)*. (F) Ridgeline diagram of enriched biological process of the significant upregulated genes. X-axis shows expression fold change of antagomir-99a-rp treated cells over negative control antagomiR. Ridgeline analysis was done with ExpressAnalyst *(68)*. (G) IgG1^+^ B cells in antagomiR-99a-5p treated normalized to negative control antagomir. *P* values were determined by two-tailed paired t-test. (H) schematic depiction of experimental setup for miRNA pull-down and downstream analysis plan for identification of miR-99a-5p direct targets based on pull-down RNAseq and bulk RNAseq. (I) Venn Diagram of upregulated genes pull-down seq ∩ upregulated genes bulk RNAseq from all 3 donors. (J) Z-score Bar graph of Genome-wide CRISPR knockout screen data in CH12F3-2 cells from *(35)*. Positive z-score indicates reduced CSR, negative z-score indicates enhanced CSR upon the knocking-out of denoted genes.

To understand the mechanism by which miR-99a-5p exerts its function and to confirm that it is B cell-dependent, we silenced miR-99a-5p in isolated B and T cells. Interestingly, treatment of B cells, but not T cells, with antagomiR-99 resulted in significant differential gene expression (Fig. 4D), thereby corroborating our previous findings of a B cell-dependent phenotype upon silencing miR-99a-5p. Hierarchical clustering heatmap highlighted clusters of genes that are associated with miR-99a-5p silencing (Fig. 4E). DEGs analysis (Log_2_FC ≥1 and q-value<0.05) revealed 155 significantly upregulated genes and 15 downregulated genes upon antagomir-99 treatment versus negative control. The 20 top upregulated genes and all downregulated genes are shown in Fig. S3A (full list in supplementary Table 1). Enrichment analysis of significantly upregulated genes highlighted the cell-cycle as one of the top pathways affected (Fig. 4F). Upregulated genes were also enriched in DNA repair signaling. Both of which are pathways involved in B cell class switch recombination (CSR), a process by which diversification of antibody isotypes is mediated. Biological process enrichment analysis (Fig. S3B), supported by upregulation of key cell cycle regulators (Fig. S3C) showed a similar trend; thereby supporting cell cycle regulation as a main arm of miR-99a-5p function. miR-99a-5p mediated gene regulation also seems to favor non-homologous end-joining (NHEJ) protein machinery over homologous recombination (HR) machinery, the former being primarily utilized by CSR. This is exhibited through the upregulation of HR key players including EME1, RAD51, and DMC1, amongst others, upon silencing of miR-99a-5p (Fig. S3D). in addition, PLXND1, which is described as a novel regulator of germinal centers and humoral immune responses *(34)*, was downregulated upon miR-99a-5p silencing in our setting. Next, we wanted to explore if miR-99a-5p indeed regulates CSR efficiency based on the aforementioned mechanisms. We treated splenic B cells with a silencer or a mimicker of miR-99a-5p with their respective controls and assessed class switching from IgM to IgG1. Strikingly, silencing miR-99a-5p resulted in a significant decrease in CSR efficiency, exhibited by lower IgG1 detection, while overexpressing miR-99a-5p with a miRNA mimic resulted in a significant increase in CSR (Fig. 4G).

### miR-99a-5p regulates B cell maturation by targeting CSR suppressors

While silencing and mimicking miR-99a-5p gave us an idea of the possible genes and pathways regulated by B cell EV miRNA signaling, it did not provide a specific downstream target mediation such function. We therefore aimed to identify miR-99a-5p direct binding targets to understand the mechanism by which it mediates its B cell phenotype using a streptavidin-bead-based pull-down assay. After treating isolated B cells with a biotinylated mimic of miR-99a-5p and biotinylated negative control, we lysed the cells and collected the miRNA-mRNA complexes by isolation with streptavidin-conjugated beads. By comparing upregulated genes in mimic-miRNA vs negative control-miRNA pulldown, we were able to identify possible targets of miR-99a-5p. Upregulated genes in the pull-down prep were then compared to upregulated genes in bulk RNAseq prep (+/-miR-99a-5p silencing) in each respective donor (Fig. 4H). The rationale is that upregulated genes upon silencing miR-99a-5p should be targets of miR-99a-5p (directly or indirectly) and by comparing them to pull-down genes we would be able to eliminate the hits that are not directly targeted by miR-99a-5p. The overlap between all comparisons resulted in 2 unique hits, namely HJURP and SOX5 (Fig. 4I). The role of these 2 genes in CSR has been independently verified in a genome-wide CRISPR knockout screen study by Feng et al *(35)* HJURP and SOX5 knockout lines showed more class-switching that wildtype lines suggesting that they are suppressor of CSR. Drug Z-scores of AICDA and TP53BP1, well-established regulating genes in CSR, in comparison to HJURP and SOX5 (Fig. 4J). HJURP and SOX5 are likely suppressing CSR through distinct but complementary mechanisms. HJURP’s role in chromatin remodeling prevents the necessary DNA regions from becoming accessible for CSR, while SOX5’s role in transcriptional regulation ensures that the regulation and fine-tuning of expression of genes required for CSR. Together, they create a robust system to maintain genomic stability, thereby preventing the necessary DNA recombination required for CSR.

### B cell isotype switching positively correlates with better melanoma prognosis

Based on our previous data, we expected that we should observe increased immunoglobulins isotype switching in ICB therapy responders over non-responders. Indeed, this was the case in biopsy samples from melanoma patients, where responders showed a significant increase in isotype class-switching (presented as IgG or IgA fold increase over IgM) in comparison to non-responders (Fig. 5 A-C). We postulate that increased class-switching enhances the anti-tumor immune response and could play a role in immune-surveillance. BCR diversification reflects a highly responsive and adaptable immune system. A diversified BCR repertoire allows for broader and more effective antigen recognition, improved antibody production, better T cell activation, and sustained immune pressure on the tumor, all of which contribute to the favorable immunotherapy response and better clinical outcomes.

**Figure 5.**
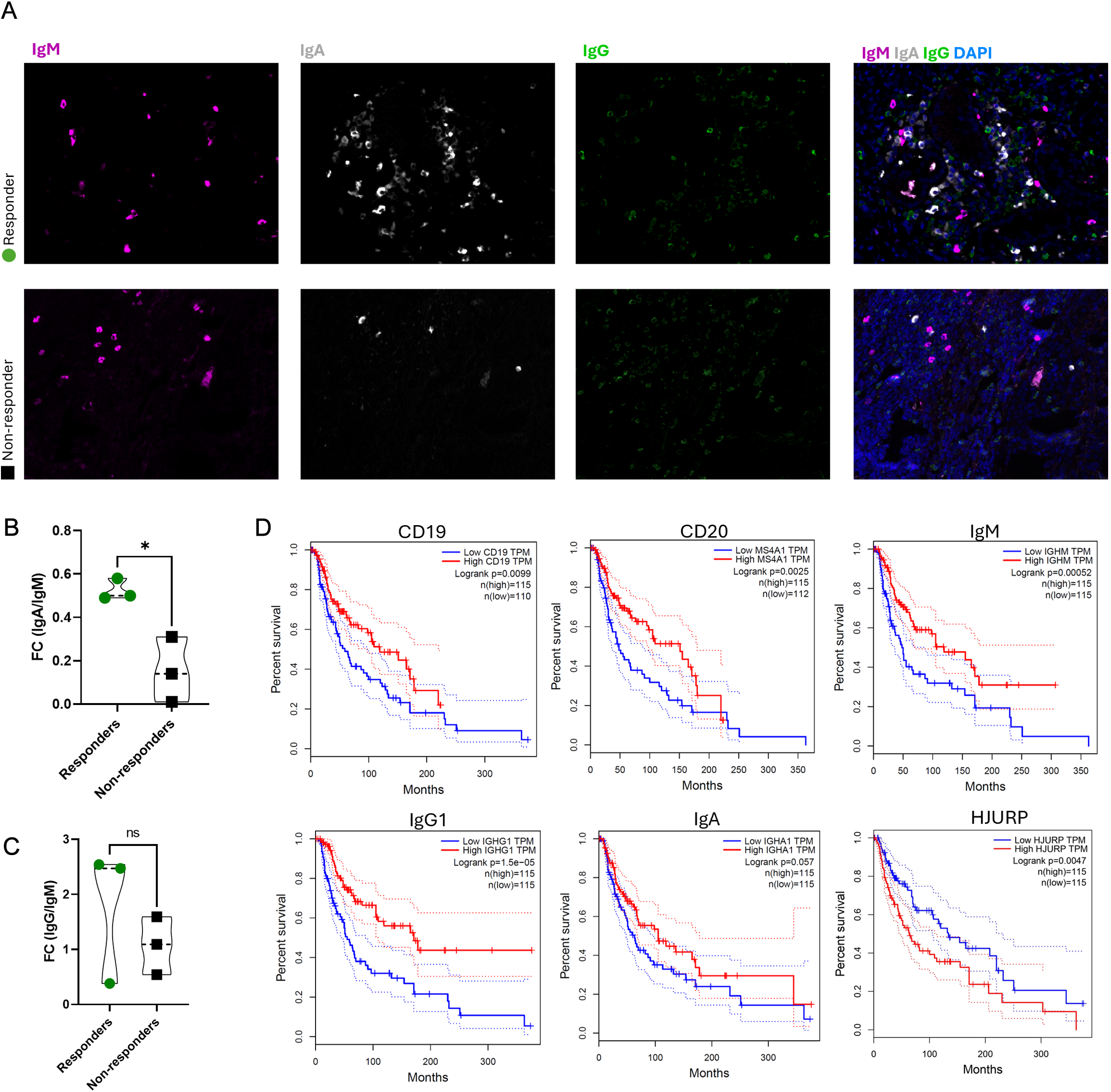
Switched Immunoglobulins are associated with increased overall survival. (A) representative images of immunofluorescence staining for IgM (magenta), IgG (green), and IgA (grey) from responders and non-responders. (B-C) quantification of immunoglobulin switching, represented as fold increase of IgA or IgG over IgM. *P* values were determined by Welch’s t-test (*p<0.05). (D) overall survival plots of melanoma patients (SKCM dataset) using GEPIA platform *(73)* (Mantel–Cox test for hypothesis test). Survival is plotted as a function of CD19, CD20, IgM, IgG1, IgA, and HJURP expression

This was in line with TCGA data, where correlation analysis was done to analyze the relationship between CD19, CD20, and immunoglobulins expression and the prognosis of melanoma by using Kaplan–Meier survival curve analysis (Fig. 5D). The results showed a significant positive correlation between the high expression of these genes and good prognosis. Whereas analysis of HJURP showed a significant positive correlation of low expression of HJURP with better survival (Fig. 5D). These findings show that i) increased presence of B cells (marked by high expression of CD19 and CD20) and enhanced antibody production (marked by high expression immunoglobulins) is correlated with better prognosis in melanoma, and ii) lower expression of HJURP, a direct target of miR99, is correlated with enhanced survival. Collectively, these findings suggest a direct role of B cell EVs in modulating the immune TME through enhancing CSR and promoting a hostile TME.

## DISCUSSION

We propose that the response to therapy relies on a specific immune cell profile at baseline, in addition to tumor-derived antigens and broad activation of T cells through ICB therapy. This conclusion can be drawn from studies showing that 1) B-cell-associated signatures were enriched in immunotherapy responders over non-responders, 2) B-cell signatures were more prominent than T-cell signatures, 3) B cell infiltration and BCR diversification are independent overall survival prognostic factors in melanoma *(36–39)*. These observations were corroborated by our meta-analysis of three independent datasets exploring transcriptome profile differences between responders and non-responders of anti-PD1 therapy at baseline *(19–21)*. We were able to show that there is a correlation between B cell tumor enrichment and response outcome. The majority of the upregulated genes in responders include many genes influencing B cell development, homing, and antibody diversification such as AID, PAX5, CXCR5/CXCL13 *(36–39)*, and that T cell signatures were far less present. Similar findings of B cells enrichment in responders’ tumor tissue were reported by Helmink et al. *(11)*. In a fourth independent scRNAseq dataset, we were able to show that the upregulated genes in responders (from the meta-analysis) are enriched in the B cell clutser, rather than in melanoma cells or any other immune compartment. Additionally, we show that these upregulated genes have distinct B cell-based functions, including B cell activation, differentiation, somatic recombination of immune receptors (immunoglobulin superfamily), and BCR signaling. Collectively these data strengthen the correlation between of B cells and ICB therapy outcome and gives support to the notion that significant change in the TME of responders happens on the B cells level.

Optimal T cell anti-tumor immune response requires B cell presence and contribution.. Reports have clearly shown depletion of B cells to decrease CD4^+^ and CD8^+^ T cells infiltration, impair their activation, and increase tumor burden *(8, 9, 40)*. In our in vitro tumor-killing assay, we were able to reproduce B cell depletion effect on impairing T cell-mediated anti-tumor activity. Interestingly, depletion of B cells did not affect the total percentage of T cell in culture, so we postulated that the impairment in T cell mediated cytotoxicity upon B cell depletion was due to noncellular B cell-based mechanism. In addition, we observed a transfer of fluorescent stain from labeled cells to non-labeled cells in our co-culture system. We were able to show that this transfer could be via the uptake of EVs from labeled cells by recipient cells. We also showed that this transfer is not passive, but deliberate, as there’s a clear differential uptake of fluorescent stain by different cell types in the co-culture system. We were also able to show that rescuing B cell-depleted cultures with B cell-derived EVs was able to rescue T cell-mediated anti-tumor activity. Collectively, this strongly supports our premise, that B cells exercise their anti-tumor role, at least in part, via EV-mediated signaling. This led us to move towards studying *in situ* EV-mediated signaling in melanoma patient tumors undergoing ICB therapy. Here, we show for the first time, the direct role of tumor in situ CD19^+^ EVs in shaping response to ICB therapy.

Our analysis of tumor biopsies of melanoma patients reproduced the same conclusion of previous reports showing an enrichment of immune cells within the tumors of responders. Strikingly, tissue-derived EVs recapitulated what we have observed thus far in the cellular TME of immune populations enrichment in responders over non-responders. This also resulted in distinct clustering of EV profiles from patients into 2 groups, that correlated with the immunotherapy response of the patients. Notably, this clustering was independent of the presence of tertiary lymphoid structures in patients’ tumors.

By comparing the miRNA cargo of B cell-derived EVs with melanoma-derived EVs, we were able to identify a set of miRNAs unique to B cell-derived EVs. The predicted gene targets of the unique B cell-derived EVs miRNA, upregulated in responders, seem to play a role in lymphocytes activation, suggesting a possible mechanism of self-regulation and activation. Interestingly, the top upregulated miRNA in responders CD19^+^ EVs was miR-99a-5p, a well-established tumor suppressor across multiple cancers *(31, 32, 41, 42)*. We propose that miR-99a-5p is selectively packaged in CD19^+^ EVs in responders, and it plays an important role in B cell-mediated anti-tumor role. Therefore, this led us to postulate that silencing miR-99a-5p could lead to impaired tumor killing in our co-culture system. Indeed, treatment with antagomiR-99a-5p was able to significantly reduce tumor killing in our co-culture system. Moreover, antagomiR-99a-5p impaired T cell mediated anti-tumor activity only in the presence of B cells, and without affecting the number of T cells in culture, further confirming miR-99a-5p B cell-dependent role.

We aimed to identify possible targets of miR-99a-5p regulation. Isolated B and T cells were treated with anitagomiR-99a-5p to silence it and sequenced for transcriptome profiling. B cells had several significant DEGs, whereas T cells had no significant DEGs, reiterating that miR-99a-5p has a B cell-dependent phenotype. Based on transcriptome data, miR-99a-5p regulates the cell cycle progression by inhibiting key cell cycle regulators, either directly or indirectly. miR-99a-5p also regulates genes involved in DNA repair and recombination, both of which are parts of the CSR process. We propose that miR-99a-5p regulates the cell cycle through prolonging G1 phase to promote the non-homologous end joining (NHEJ) repair pathway and subsequently promote CSR. This notion could be partially supported by the upregulation of E2F family genes, CDC25A, CDK1, and Cyclin E2 (CCNE2), amongst other factors known to regulate G1/S phase transition, upon miR-99a-5p silencing *(43–46)*. This has been independently verified in RCC cells, where expressing miR-99a-5p exogenously induced G1-phase cell cycle arrest through targeting several cyclins in RCC cells *(47)*. CSR is shown to be more active in G1 to early S phase as DNA is more accessible and DNA repair factors in CSR are cell cycle-dependent *(48, 49)* miR-99a-5p might also favor NHEJ over HR through the downregulation of important factors in the process such as FOXM1, RAD51, RAD54L, and NEIL3 amongst others*(50–55)*. In addition, PLXND1 was downregulated upon miR-99a-5p silencing in our setting. A study has reported that PLXND1(-/-) mice displayed defective GC reactions in T cell-dependent immune activation and defective production of IgG1 and IgG2b *(56)*. Collectively, these conclusions were validated by miR-99a-5p involvement in promoting CSR in ex-vivo splenic B cells. In our hands, silencing miR-99a-5p resulted in a significant decrease in CSR exhibited by lower IgG1 detection in antagomir-99a-5p treated cells. In contrast, overexpressing miR-99a-5p resulted in a significant increase in CSR in mimicker-99a-5p treated cells.

Our miRNA pull-down assay identified 2 targets of miR-99a-5p in B cells, HJURP and SOX5. Both of which have been independently shown to negatively affect CSR (50). HJURP plays a role in chromatin remodeling and is involved in the DDR pathway through promotion of HR repair pathway *(57)*. SOX5, as a transcription factor, regulates gene expression of select genes, and has been shown to regulate B cells proliferation and differentiation into plasmablasts, a short-lived lower-affinity antibody producing cells *(58)*. Hence, HJURP and SOX5 are likely suppressing CSR through distinct but complementary mechanisms. These findings go hand in hand with the conclusions we drew from all the earlier data showcasing miR-99a-5p role in regulating the cell cycle and DDR pathways, resulting in enhanced CSR.

Our data suggest that B cells EVs-derived miR-99a-5p exerts its function in promoting a positive immunotherapy response possibly via two arms, 1) the regulation of the cell cycle and DNA repair pathways in B cells; and subsequently 2) through CSR promotion to induce a diverse antibody repertoire. Switched B cells were reported to be enriched in the tumors of immunotherapy responders with high clonal expansion *(59, 60)*. Increased BCR class switching and affinity maturation, and BCR variable region diversification in responders were identified as an independent overall survival prognostic factor in melanoma *(59, 60)*; suggesting B cells actively participate in shaping anti-tumor immunity. These observations were validated in our hands, where ICB therapy responders showed a significant increase in isotype class-switching in comparison to non-responders. These observations were additionally validated with our overall survival analysis based on CD19, CD20, IgM, IgG, IgA expression in melanoma TCGA dataset. High expression of CD19 and CD20 is associated with significantly better overall survival. Nonetheless, what was striking is that immunoglobulins had a more declared positive outcome, with IgG (presenting a switched phenotype) exhibiting higher correlation than IgM with overall survival. In addition, high expression of HJURP has been linked to poor overall survival in different cancers *(61)*. As we have shown, HJURP is a direct target of miR-99a-5p and that its inhibition results in enhanced CSR *(35)*. In melanoma, lower expression of HJURP is correlated with significantly better overall survival (Fig. 5D). These findings suggest that B cells-derived EVs core function is to regulate B cells themselves, at least in the case of miR-99a-5p, to promote B cell maturation, antibody diversification, and an anti-tumor B cell population. Interestingly, several exciting reports on B cell-derived EVs show that they can carry p-MHC complexes *(62)*, functional BCRs *(63)*, and functional IgM *(64)* and IgG *(65)*, which in turn can be uptaken by recipient cells to modulate their functions. This strongly suggests a possible role of B cell-derived EVs in immunosurveillance through antigen presentation and B cell maturation. B cell maturation can improve their tumor antigen uptake and presentation as BCRs. Subsequently, this B cell-mediated anti-tumor profile can shape the TME at large, and T cells specifically, to mediate a hostile environment to target and eradicate the tumor. This supports the notion that B cells might be as essential as T cells in shaping immunotherapy outcome. With this understanding, B cells potential can be taken into consideration when designing anticancer therapies to fully harness what B cells have to offer. In addition, identifying a key-regulator of B cell response in responders to immunotherapy, such as miR-99a-5p, could revolutionize the way we think of immunotherapy with more focus on the entirety of the immune TME, and less on T cells as the final player in the immune response.

## MATERIALS AND METHODS

### Patient samples and tissue processing

Melanoma slow-frozen patient biopsies were obtained from the biobank of the Dermatology Department of the University Hospital Zürich (KEK_PB_2018-00194). All samples used were surplus materials from routine surgeries. Collection, preparation, and freezing of the fresh biopsies are outlined in *(66)*. Informed consent had been obtained from all patients and all experiments conducted according to the ethical rules of the Cantonal Ethic Committee of Zurich (Ethics form BASEC: 2014-0425). Tissue processing protocol is detailed in (*(27)*. Briefly, cryovials were thawed in 37°C water bath and tissue fragments were collected into a pre-cooled 35mm dish. Tissue fragments were washed and sliced into smaller pieces in 1 mL of digestion enzyme mixture of Dispase-II (5mg/mL final concentration) and DNase-I (10ug/mL final concentration). Tissue fragments and digestion supernatant were collected into 2 mL eppendorf tube and incubated at 37°C with shaking in a heat-block for 2 hrs. Samples were centrifuged at 500xg for 5 mins to clear EVs from cells and other debris. Supernatant containing the EVs was collected into a fresh tube and kept at 4°C until analysis.

### Tissue biopsy flow cytometry

Tissue fragments were further processed by incubating them with 1 mL of Collagenase-IA (0.5 mg/mL final concentration) at 37°C with shaking in a heat-block for 45 mins. Samples were centrifuged at 500 x g for 5 mins to separate cells from EVs. After centrifugation, cell suspension was passed through 70uM pre-wetted cell strainer. With the back of 5 mL syringe plunger, remaining tissue was mildly dissociated, and an additional 5 ml of media was used to wash the cells through. Single cell suspension was then stained for FACS analysis and stained with Live/Dead blue (Invitrogen, L23105) and antibodies against CD45 (Biolegend, 304012), CD19 (Biolegend, 302233), CD3 (Biolegend, 300305). Samples were acquired using Cytek Aurora 5L.

### EVs isolation and characterization

Digestion supernatant was serially centrifuged to clear from cells and debris. Supernatant was centrifuged for 10 minutes at 300 x g and 2000 x g. Supernatant was then centrifuged at 10,000 x g for 30 mins before ultracentrifugation at 100,000 X g for 1 hour at 4°C. After ultracentrifugation, EVs pellet was resuspended in 0.22 µm filtered PBS for downstream analysis. EV characterization was carried out using transmission electron microscopy (TEM) and nano-flow cytometry. With TEM, we were able to visualize the morphology and size of EVs isolated from processed tissue samples. For TEM, EVs samples were transferred onto pioloform-coated EM copper grids by floating the grids on a droplet containing freshly prepared EVs suspension placed on parafilm and incubated for 5 mins. The grids were washed 3 times for 5 mins each before contrasting bound EVs with a mixture of 2% w/v methyl cellulose and 2% w/v uranyl acetate (9:1 ratio) on ice for 10 mins. Grids were then allowed to air-dry before imaging. TEM images were analyzed with MAPS software (Thermo Fisher) for EVs sizes estimation. The same samples were sized with nano flow-cytometry with nFCM sizing beads made of a 68 nm, 91 nm, 113 nm, and 155 nm beads cocktail. Both size estimations were compared by grouping TEM sized particles into the 4 size categories of nFCM beads (0-75 nm sized particles are assigned to 68 nm category, 76-98 nm to 91 nm category, 99 - 130 nm to 113 nm category, 131-180 nm to 155 nm category).

### EVs surface marker analysis

We aimed at characterizing surface and cargo molecules of tissue-derived EVs from responders and non-responders melanoma patients undergoing ICB therapy. To characterize EVs surface proteins, we utilized a multiplex beads assay for the detection of 37 broad markers simultaneously, including several immune markers to provide an overview of the EVs landscape. Briefly, 120 uL of resuspended EVs samples were incubated with 15 uL EVs capture beads overnight at room temperature in Thermomixer orbital shaker (Eppendorf). EVs and bead mixture was then washed with 500 uL MACSPlex buffer and centrifuged at 3000xg for 5 minutes at room temperature. EVs-beads conjugates were then incubated with 15 uL EVs detection reagent (5 uL of CD9, CD63, CD81 detection reagents) for 1 hour at room temperature in Thermomixer orbital shaker (Eppendorf). Samples were washed with 500 uL MACSPlex buffer and centrifuged at 3000xg for 5 minutes at room temperature. 500 uL of supernatant was discarded and the rest of the samples were resuspended and transferred into FACS tubes for downstream analysis. Blank controls (beads and Macsplex buffer) were used to deduct background signal (Fig. S2). Samples were analyzed using LSRII Fortessa 4L FACS machine. For immunostaining EVs, isolated EVs samples were stained in 100 uL of antibody staining solution at a dilution of 1:100 for 1 hour on ice. Antibodies against melanoma (melanoma cocktail, Novus, NBP2-34547B), CD63 (Biolegend, 353005), CD45 (Biolegend, 304012), were used. Additionally, EVs were stained with cell trace (1:1000, C34564). Excess antibodies and cell trace were washed out to reduce background signal by topping up samples with 1 mL 0.22 um filtered PBS before ultracentrifugation at 100,000 X g for 1 hour at 4°C. Supernatant was discarded and the EV pellets were resuspended in 50 uL of 0.22 um filtered PBS for downstream analysis.

Experimental controls included PBS blank controls, antibody staining solution without EVs, and EVs stained with isotype controls. These controls were prepared to exclude any background signal or non-specific binding of antibodies (Fig. S2). The nanoanalyzer alignment was adjusted in every experimental setup using 250 nm QC nanosphere beads at a concentration of 2.07×1010 particles/mL (Lot 2012141, nFCM). Silica sizing beads (S16M-Exo, nanoFCM) composed of 4 beads populations of 68, 91, 113, and 155 nm sizes were used to estimate the size of recorded particles. All beads were diluted at 1:100 for machine setup. Laser power was kept at 20 mW and 40 mW for the 488 nm and 640 nm lasers, respectively. Sampling pressure was kept at 0.8 kPa, and samples recorded were kept at 2000-12,000 events/minute. Concentration and size estimation of all samples were obtained using NF profession V2.18 software and downstream analysis was carried out using Flowjo software.

### PBMCs Bulk RNA sequencing

PBMCs from healthy donors were thawed, washed twice with media, and resuspended in wash buffer (2% FBS 1mM EDTA PBS). Cell suspension was separated into 2 tubes for B and T cells isolation using StemCell EasySep™ Human CD19 Positive Selection Sample Kit II or EasySep™ Release Human CD3 Positive Selection Kit, respectively. Isolated B and T cells were seeded in 48-well plates and allowed to rest for 2 hours before stimulation with SEB (50ng/mL) for 4 hours. Isolated cells were treated with antagomir-99a-5p or negative control at a final concentration of 300 nM for 48 hours. Cells wee collected, washed, and lysed with trizol prior to total RNA isolation with RNeasy Mini Kit (Qiagen, Germany). The concentration and purity of total RNA were assessed using NanoDrop2000. Quality control of RNA samples was performed with Agilent 2100 Bioanalyzer RNA 6000 Nano Kit. Library preparation and sequencing was done with DNBSEQ™ platform (BGI Genomics). DEGs analysis was done using DESeq2 on Dr.Tom platform (Beijing Genomics Institute, BGI). Cutoff for significant genes was set at Log_2_FC ≥1, and q-value<0.05. Ridgeline analysis of enriched biological process was carried out for the significant upregulated genes. X-axis shows expression fold change of antagomir-99a-rp treated cells over negative control antagomiR. Ridgeline analysis was done with ExpressAnalyst *(68) Class Switch recombination assay* Splenic primary B cells were cultured in RPMI 1640 media with 20% FCS (Sigma Aldrich), 5% Medium NCTC-109 (Thermo Fisher), 1X Penicillin-Streptomycin-Glutamine (Gibco), 1mM Sodium Pyruvate (Gibco), 1X MEM-NEAA (Gibco), 2mM GlutaMAX (Gibco), and 55µM β-mercaptoethanol (Pan Biotech). To induce CSR from IgM to IgG, primary B cells were activated with LPS (25 ug/mL) and IL-4 (10 ng/mL). Cells were treated with 1 uM final concentration of miR-99a-5p silencer and mimicker, in addition to their respective negative controls. Cells were incubated for 96 hrs. At termination points, cells were collected, washed 2 times with cold FACS buffer with centrifugation at 400xg for 4 minutes at 4C. Cells were stained with anti-Mo IgG (Invitrogen), anti-Mo IgM (Invitrogen) and Zombie Violet™ Fixable Viability (Biolegend) for 30 minutes. Cells were washed two additional times and centrifuged before resuspension in PBS. Samples were acquired using Cytek Aurora 5L.

### Immunostaining and quantitative pathology

Immunophenotyping was performed using a six-colour multispectral immunofluorescence protocol. Briefly, 3 µm thick FFPE tissue sections were mounted on HistoBond adhesive microscope slides (Marienfeld, Germany). Slides were baked at 70 C for 5 hours, deparaffinized and rehydrated through serial ethanol concentrations. Next, they were placed in Tris-EDTA pH 9.0 buffer and heated at +95°C for 80 minutes for antigen retrieval, then blocked using 4% BSA/0.01% Triton X-100/TBS. Antibodies against CD45 (Dako, M0701), IgG (Abcam, ab307524), IgM (Dako, A0425), and IgA (Southern Biotech, 2050-07) were used and diluted in 1% BSA/0.01% Triton X-100/TBS. After overnight incubation with primary antibodies at 4 C, the slides were rinsed and the tissue sections were incubated with secondary antibodies labelled with fluorophores (JacksonImmunoresearch) for 1 hour at room temperature. Finally, the slides were incubated with DAPI (Thermo Fisher Scientific) for 5 minutes at room temperature and then mounted using VECTASHIEL PLUS Antifade mounting medium (Vector Labs). Whole slides were imaged using PhenoImager HT 2.0 (Akoya), a multispectral image scanner and multispectral images of 20x high power field were captured. The images were subjected to spectral unmixing techniques using Inform v. 2.7 (Akoya). The image data files were then converted into flow cytometry data format using Biobase and Flowcore (R studio), and further analysis to identify and quantify distinct cell populations was performed through Flowjo v.10.10.0 software.

### Processing of bulk RNA-seq melanoma datasets

Bulk RNA-seq melanoma datasets from Hossain et al. *(19)*,Hugo et al. *(20)*, and Riaz et al. *(21)* were downloaded from NCBI Gene Expression Omnibus (GEO) under the accession numbers GSE213145, GSE78220 and GSE91061, respectively. Raw sequencing fastq files from each of the three datasets were downloaded using the package sratools (v 3.0.3) and their primary quality control was performed using the software FastQC (v 0.11.9).Next, in each of the files, the package Cutadapt (v 1.18) was utilized to trim the adapters, filter low-quality bases from the 3’-end of the reads (average base quality threshold > 15), and remove reads shorter than 35 base pairs and with more than 20 unidentified bases. Following this step, the secondary quality control of the files was carried out to validate Cutadapt results. The reads were then mapped to the human GENCODE GRCh38.p13 reference genome using the splice-aware aligner GSNAP (v 2021-08-25) ensued by the implementation of package RSeQC (v 5.0.1) for the file mapping quality check. Mapping quality control comprised the verification of mapped read distribution across different genome regions and the percentage quantification of ribosomal RNA contaminants. Packages MultiQC (v 1.14) and Samtools (v 1.10) were used in summarizing the file reports from each step and manipulating the sam/bam files, respectively. Finally, the table of the raw read counts was generated through the function featureCounts (v 2.0.3) for each of the samples passing the quality controls. Differential gene expression analysis of melanoma datasets was performed using the DESeq2 (v 1.38.0) package in R (v 4.2.3). Batch correction of the read counts was implemented with the ComBat function in sva (v 3.48.0) package. Immune infiltration estimation was carried out using immune deconvolution method TIMER2.0 *(69)* based on bulk RNAseq data. Protein-protein interaction network analysis was done using NetworkAnalyst *(70)*. IMEx Interactome was selected as reference database.

## AUTHORS CONTRIBUTION

AA and RC conceived the study and designed the experiments. AA conducted the study and wrote the manuscript. RC provided funding, conceptual insights, discussions, and edited the manuscript. AB generated the meta-analysis dataset of ICB patient and provided bioinformatics support. KCG conducted the CSR experiment. KG and KS performed multispectral immunofluorescence imaging and analysis. SB analyzed biopsy flow cytometry data and provided insights. MPL and JMMG provided patient samples and insights.

## ACKNOWLEDGMENTS

We thank all members of the melanoma biobank at USZ for their help and feedback; UZH flow cytometry facility for their continuous support; Johannes Riemann and UZH microscopy facility for TEM analysis. Prof C Münz, I Arnold, and O Chijoke for critical reading of the manuscript.

## REFERENCES

1. J. Galon, B. Mlecnik, G. Bindea, H. K. Angell, A. Berger, C. Lagorce, A. Lugli, I. Zlobec, A. Hartmann, C. Bifulco, I. D. Nagtegaal, R. Palmqvist, G. V Masucci, G. Botti, F. Tatangelo, P. Delrio, M. Maio, L. Laghi, F. Grizzi, M. Asslaber, C. D’Arrigo, F. Vidal Vanaclocha, E. Zavadova, L. Chouchane, P. S. Ohashi, S. Hafezi Bakhtiari, B. G. Wouters, M. Roehrl, L. Nguyen, Y. Kawakami, S. Hazama, K. Okuno, S. Ogino, P. Gibbs, P. Waring, N. Sato, T. Torigoe, K. Itoh, P. S. Patel, S. N. Shukla, Y. Wang, S. Kopetz, F. A. Sinicrope, V. Scripcariu, P. A. Ascierto, F. M. Marincola, B. A. Fox, F. Pagès, Towards the introduction of the ‘Immunoscore’ in the classification of malignant tumours. J Pathol 232, 199–209 (2014).

2. W. H. Fridman, F. Pagès, C. Sautès-Fridman, J. Galon, The immune contexture in human tumours: impact on clinical outcome. Nat Rev Cancer 12, 298–306 (2012).

3. F. Maibach, H. Sadozai, S. M. Seyed Jafari, R. E. Hunger, M. Schenk, Tumor-Infiltrating Lymphocytes and Their Prognostic Value in Cutaneous Melanoma. Front Immunol 11 (2020), doi:10.3389/fimmu.2020.02105.

4. Q. Li, S. Teitz-Tennenbaum, E. J. Donald, M. Li, A. E. Chang, In Vivo Sensitized and In Vitro Activated B Cells Mediate Tumor Regression in Cancer Adoptive Immunotherapy. The Journal of Immunology (2009), doi:10.4049/jimmunol.0803773.

5. F. Martin, A. C. Chan, B CELL IMMUNOBIOLOGY IN DISEASE: Evolving Concepts from the Clinic. Annu Rev Immunol (2006), doi:10.1146/annurev.immunol.24.021605.090517.

6. B. H. Nelson, CD20+ B Cells: The Other Tumor-Infiltrating Lymphocytes. The Journal of Immunology 185, 4977–4982 (2010).

7. C. B. Rodgers, C. J. Mustard, R. T. McLean, S. Hutchison, A. L. Pritchard, A B-cell or a key player? The different roles of B-cells and antibodies in melanoma. Pigment Cell Melanoma Res 35, 303–319 (2022).

8. J. Griss, W. Bauer, C. Wagner, M. Simon, M. Chen, K. Grabmeier-Pfistershammer, M. Maurer-Granofszky, F. Roka, T. Penz, C. Bock, G. Zhang, M. Herlyn, K. Glatz, H. Läubli, K. D. Mertz, P. Petzelbauer, T. Wiesner, M. Hartl, W. F. Pickl, R. Somasundaram, P. Steinberger, S. N. Wagner, B cells sustain inflammation and predict response to immune checkpoint blockade in human melanoma. Nature Communications 2019 10:*1* 10, 1–14 (2019).

9. D. J. DiLillo, K. Yanaba, T. F. Tedder, B Cells Are Required for Optimal CD4+ and CD8+ T Cell Tumor Immunity: Therapeutic B Cell Depletion Enhances B16 Melanoma Growth in Mice. The Journal of Immunology 184, 4006–4016 (2010).

10. R. Cabrita, M. Lauss, A. Sanna, M. Donia, M. Skaarup Larsen, S. Mitra, I. Johansson, B. Phung, K. Harbst, J. Vallon-Christersson, A. van Schoiack, K. Lövgren, S. Warren, K. Jirström, H. Olsson, K. Pietras, C. Ingvar, K. Isaksson, D. Schadendorf, H. Schmidt, L. Bastholt, A. Carneiro, J. A. Wargo, I. M. Svane, G. Jönsson, Tertiary lymphoid structures improve immunotherapy and survival in melanoma. Nature 577 (2020), doi:10.1038/s41586-019-1914-8.

11. A. B. A. Helmink, S. M. Reddy, J. Gao, S. Zhang, R. Basar, R. Thakur, K. Yizhak, M. Sade-Feldman, J. Blando, G. Han, V. Gopalakrishnan, Y. Xi, H. Zhao, R. N. Amaria, H. A. Tawbi, P. Cogdill, W. Liu, V. S. LeBleu, F. G. Kugeratski, S. Patel, M. A. Davies, P. Hwu, J. E. Lee, J. E. Gershenwald, A. Lucci, R. Arora, S. Woodman, E. Z. Keung, P. O. Gaudreau, A. Reuben, C. N. Spencer, E. M. Burton, L. E. Haydu, A. J. Lazar, R. Zapassodi, C. W. Hudgens, D. A. Ledesma, S. F. Ong, M. Bailey, S. Warren, D. Rao, O. Krijgsman, E. A. Rozeman, D. Peeper, C. U. Blank, T. N. Schumacher, L. H. Butterfield, M. A. Zelazowska, K. M. McBride, R. Kalluri, J. Allison, F. Petitprez, W. H. Fridman, C. Sautès-Fridman, N. Hacohen, K. Rezvani, P. Sharma, M. T. Tetzlaff, L. Wang, J. A. Wargo, B cells and tertiary lymphoid structures promote immunotherapy response. Nature 2020 577:7791 577, 549–555 (2020).

11. G. Chiaruttini, S. Mele, J. Opzoomer, S. Crescioli, K. M. Ilieva, K. E. Lacy, S. N. Karagiannis, B cells and the humoral response in melanoma: The overlooked players of the tumor microenvironment. Oncoimmunology 6, 1–11 (2017).

12. S. Yu, H. Cao, B. Shen, J. Feng, Tumor-derived exosomes in cancer progression and treatment failure. Oncotarget 6, 37151–37168 (2015).

13. G. Chen, A. C. Huang, W. Zhang, G. Zhang, M. Wu, W. Xu, Z. Yu, J. Yang, B. Wang, H. Sun, H. Xia, Q. Man, W. Zhong, L. F. Antelo, B. Wu, X. Xiong, X. Liu, L. Guan, T. Li, S. Liu, R. Yang, Y. Lu, L. Dong, S. McGettigan, R. Somasundaram, R. Radhakrishnan, G. Mills, Y. Lu, J. Kim, Y. H. Chen, H. Dong, Y. Zhao, G. C. Karakousis, T. C. Mitchell, L. M. Schuchter, M. Herlyn, E. J. Wherry, X. Xu, W. Guo, Exosomal PD-L1 contributes to immunosuppression and is associated with anti-PD-1 response. Nature 560, 382–386 (2018).

14. M. Poggio, T. Hu, C. C. Pai, B. Chu, C. D. Belair, A. Chang, E. Montabana, U. E. Lang, Q. Fu, L. Fong, R. Blelloch, Suppression of Exosomal PD-L1 Induces Systemic Anti-tumor Immunity and Memory. Cell 177, 414–427.e13 (2019).

15. F. Xie, M. Xu, J. Lu, L. Mao, S. Wang, The role of exosomal PD-L1 in tumor progression and immunotherapy. Mol Cancer 18, 146 (2019).

16. V. R. Juneja, K. A. McGuire, R. T. Manguso, M. W. LaFleur, N. Collins, W. N. Haining, G. J. Freeman, A. H. Sharpe, PD-L1 on tumor cells is sufficient for immune evasion in immunogenic tumors and inhibits CD8 T cell cytotoxicity. Journal of Experimental Medicine 214, 895–904 (2017).

17. N. Chaput, C. Théry, Exosomes: Immune properties and potential clinical implementations Semin Immunopathol (2011), doi:10.1007/s00281-010-0233-9.

18. S. M. Hossain, G. Gimenez, P. A. Stockwell, P. Tsai, C. G. Print, J. Rys, B. Cybulska-Stopa, M. Ratajska, A. Harazin-Lechowska, S. Almomani, C. Jackson, A. Chatterjee, M. R. Eccles, Innate immune checkpoint inhibitor resistance is associated with melanoma sub-types exhibiting invasive and de-differentiated gene expression signatures. Front Immunol 13 (2022), doi:10.3389/fimmu.2022.955063.

19. W. Hugo, J. M. Zaretsky, L. Sun, C. Song, B. H. Moreno, S. Hu-Lieskovan, B. Berent-Maoz, J. Pang, B. Chmielowski, G. Cherry, E. Seja, S. Lomeli, X. Kong, M. C. Kelley, J. A. Sosman, D. B. Johnson, A. Ribas, R. S. Lo, Genomic and Transcriptomic Features of Response to Anti-PD-1 Therapy in Metastatic Melanoma. Cell 165, 35–44 (2016).

20. N. Riaz, J. J. Havel, V. Makarov, A. Desrichard, W. J. Urba, J. S. Sims, F. S. Hodi, S. Martín-Algarra, R. Mandal, W. H. Sharfman, S. Bhatia, W. J. Hwu, T. F. Gajewski, C. L. Slingluff, D. Chowell, S. M. Kendall, H. Chang, R. Shah, F. Kuo, L. G. T. Morris, J. W. Sidhom, J. P. Schneck, C. E. Horak, N. Weinhold, T. A. Chan, Tumor and Microenvironment Evolution during Immunotherapy with Nivolumab. Cell 171, 934–949.e16 (2017).

21. L. Jerby-Arnon, P. Shah, M. S. Cuoco, C. Rodman, M. J. Su, J. C. Melms, R. Leeson, A. Kanodia, S. Mei, J. R. Lin, S. Wang, B. Rabasha, D. Liu, G. Zhang, C. Margolais, O. Ashenberg, P. A. Ott, E. I. Buchbinder, R. Haq, F. S. Hodi, G. M. Boland, R. J. Sullivan, D. T. Frederick, B. Miao, T. Moll, K. T. Flaherty, M. Herlyn, R. W. Jenkins, R. Thummalapalli, M. S. Kowalczyk, I. Cañadas, B. Schilling, A. N. R. Cartwright, A. M. Luoma, S. Malu, P. Hwu, C. Bernatchez, M. A. Forget, D. A. Barbie, A. K. Shalek, I. Tirosh, P. K. Sorger, K. Wucherpfennig, E. M. Van Allen, D. Schadendorf, B. E. Johnson, A. Rotem, O. Rozenblatt-Rosen, L. A. Garraway, C. H. Yoon, B. Izar, A. Regev, A Cancer Cell Program Promotes T Cell Exclusion and Resistance to Checkpoint Blockade. Cell 175, 984–997.e24 (2018).

22. J. D. McLeod, L. S. K. Walker, Y. I. Patel, G. Boulougouris, D. M. Sansom, Activation of Human T Cells with Superantigen (Staphylococcal Enterotoxin B) and CD28 Confers Resistance to Apoptosis via CD95. The Journal of Immunology 160, 2072–2079 (1998).

23. G. Seprényi, T. Shibata, R. Ónody, T. Kohsaka, In staphylococcus enterotoxin B (SEB)-stimulated human PBMC, the LAK activity of non-T cells might have a major role in the mechanism of glomerular endothelial cells’ injury. Immunobiology 197, 44–54 (1997).

24. J. Banér, P. Marits, M. Nilsson, O. Winqvist, U. Landegren, Analysis of T-Cell Receptor Vβ Gene Repertoires after Immune Stimulation and in Malignancy by Use of Padlock Probes and Microarrays. Clin Chem 51, 768–775 (2005).

25. K. H. W. Yim, A. Al Hrout, S. Borgoni, R. Chahwan, Extracellular vesicles orchestrate immune and tumor interaction networks Cancers (Basel) 12, 1–23 (2020).

26. A. Al Hrout, M. P. Levesque, R. Chahwan, Investigating the tumor-immune microenvironment through extracellular vesicles from frozen patient biopsies and 3D cultures. Front Immunol 14 (2023), doi:10.3389/fimmu.2023.1176175.

27. K. Ho, W. Yim, S. Borgoni, R. Chahwan, Serum extracellular vesicles profiling is associated with COVID-19 progression and immune responses. Journal of Extracellular Biology 1, e37 (2022).

28. K. H. W. Yim, O. Krzyzaniak, A. Al Hrout, B. Peacock, R. Chahwan, Assessing Extracellular Vesicles in Human Biofluids Using Flow-Based Analyzers. Adv Healthc Mater 12, 2301706 (2023).

29. J. Zhang, H. Jin, H. Liu, S. Iv, B. Wang, R. Wang, H. Liu, M. Ding, Y. Yang, L. Li, J. Zhang, S. Fu, D. Xie, M. Wu, W. Zhou, Q. Qian, MiRNA-99a directly regulates AGO2 through translational repression in hepatocellular carcinoma. Oncogenesis 2014 3:*4* 3, e97–e97 (2014).

30. T. F. Tsai, J. F. Lin, K. Y. Chou, Y. C. Lin, H. E. Chen, T. I. S. Hwang, miR-99a-5p acts as tumor suppressor via targeting to mTOR and enhances RAD001-induced apoptosis in human urinary bladder urothelial carcinoma cells. Onco Targets Ther 11, 239 (2018).

31. X. Long, Y. Shi, P. Ye, J. Guo, Q. Zhou, Y. Tang, MicroRNA-99a Suppresses Breast Cancer Progression by Targeting FGFR3. Front Oncol 9, 499482 (2020).

32. Z. Wang, Z. Zhao, Y. Yang, M. Luo, M. Zhang, X. Wang, L. Liu, N. Hou, Q. Guo, T. Song, B. Guo, C. Huang, MiR-99b-5p and miR-203a-3p Function as Tumor Suppressors by Targeting IGF-1R in Gastric Cancer. Scientific Reports 2018 8:*1* 8, 1–12 (2018).

33. E. K. Holl, B. P. O’Connor, T. M. Holl, K. E. Roney, A. G. Zimmermann, S. Jha, G. Kelsoe, J. P.-Y. Ting, Plexin-D1 Is a Novel Regulator of Germinal Centers and Humoral Immune Responses. The Journal of Immunology 186, 5603–5611 (2011).

34. Y. Feng, C. Li, J. A. Stewart, P. Barbulescu, N. S. Desivo, A. Álvarez-Quilón, R. C. Pezo, M. L. W. Perera, K. Chan, A. H. Y. Tong, R. Mohamad-Ramshan, M. Berru, D. Nakib, G. Li, G. A. Kardar, J. R. Carlyle, J. Moffat, D. Durocher, J. M. Di Noia, A. S. Bhagwat, A. Martin, FAM72A antagonizes UNG2 to promote mutagenic repair during antibody maturation. Nature *2021* 600:7888 600, 324–328 (2021).

35. M. Fuxa, M. Busslinger, Reporter Gene Insertions Reveal a Strictly B Lymphoid-Specific Expression Pattern of *Pax5* in Support of Its B Cell Identity Function. The Journal of Immunology 178, 3031–3037 (2007).

36. C. Cobaleda, A. Schebesta, A. Delogu, M. Busslinger, Pax5: the guardian of B cell identity and function. Nat Immunol 8, 463–470 (2007).

37. L. Calderón, K. Schindler, S. G. Malin, A. Schebesta, Q. Sun, T. Schwickert, C. Alberti, M. Fischer, M. Jaritz, H. Tagoh, A. Ebert, M. Minnich, A. Liston, L. Cochella, M. Busslinger, Pax5 regulates B cell immunity by promoting PI3K signaling via PTEN down-regulation. Sci Immunol 6 (2021), doi:10.1126/sciimmunol.abg5003.

38. K. M. Ansel, V. N. Ngo, P. L. Hyman, S. A. Luther, R. Förster, J. D. Sedgwick, J. L. Browning, M. Lipp, J. G. Cyster, A chemokine-driven positive feedback loop organizes lymphoid follicles. Nature 406, 309–314 (2000).

39. S. Singh, J. Roszik, N. Saini, V. K. Singh, K. Bavisi, Z. Wang, L. T. Vien, Z. Yang, S. Kundu, R. E. Davis, L. Bover, A. Diab, S. S. Neelapu, W. W. Overwijk, K. Rai, M. Singh, B Cells Are Required to Generate Optimal Anti-Melanoma Immunity in Response to Checkpoint Blockade. Front Immunol 13, 794684 (2022).

40. X. J. Li, X. Q. Luo, B. W. Han, F. T. Duan, P. P. Wei, Y. Q. Chen, MicroRNA-100/99a, deregulated in acute lymphoblastic leukaemia, suppress proliferation and promote apoptosis by regulating the FKBP51 and IGF1R/mTOR signalling pathways. British Journal of Cancer 2013 109:*8* 109, 2189–2198 (2013).

41. A. Feliciano, Y. Garcia-Mayea, L. Jubierre, C. Mir, M. Hummel, J. Castellvi, J. Hernández-Losa, R. Paciucci, I. Sansano, Y. Sun, S. Ramón Y Cajal, H. Kondon, A. Soriano, M. Segura, A. Lyakhovich, M. E. Lleonart, miR-99a reveals two novel oncogenic proteins E2F2 and EMR2 and represses stemness in lung cancer. Cell Death & Disease 2017 8:*10* 8, e3141–e3141 (2017).

42. H.-Z. Chen, S.-Y. Tsai, G. Leone, Emerging roles of E2Fs in cancer: an exit from cell cycle control. Nat Rev Cancer 9, 785–797 (2009).

43. T. Shen, S. Huang, The Role of Cdc25A in the Regulation of Cell Proliferation and Apoptosis. Anticancer Agents Med Chem 12, 631–639 (2012).

44. J. M. Enserink, R. D. Kolodner, An overview of Cdk1-controlled targets and processes. Cell Div 5, 11 (2010).

45. S. Mazumder, B. Gong, A. Almasan, Cyclin E induction by genotoxic stress leads to apoptosis of hematopoietic cells. Oncogene 19, 2828–2835 (2000).

46. L. Cui, H. Zhou, H. Zhao, Y. Zhou, R. Xu, X. Xu, L. Zheng, Z. Xue, W. Xia, B. Zhang, T. Ding, Y. Cao, Z. Tian, Q. Shi, X. He, MicroRNA-99a induces G1-phase cell cycle arrest and suppresses tumorigenicity in renal cell carcinoma. BMC Cancer 12, 1–11 (2012).

47. S. Petersen, R. Casellas, B. Reina-San-Martin, H. T. Chen, M. J. Difilippantonio, P. C. Wilson, L. Hanitsch, A. Celeste, M. Muramatsu, D. R. Pilch, C. Redon, T. Ried, W. M. Bonner, T. Honjo, M. C. Nussenzweig, A. Nussenzweig, AID is required to initiate Nbs1/γ-H2AX focus formation and mutations at sites of class switching. Nature 2001 414:6864 414, 660–665 (2001).

48. C. Rada, G. T. Williams, H. Nilsen, D. E. Barnes, T. Lindahl, M. S. Neuberger, Immunoglobulin Isotype Switching Is Inhibited and Somatic Hypermutation Perturbed in UNG-Deficient Mice. Current Biology 12, 1748–1755 (2002).

49. N. Li, Y. Xu, H. Chen, L. Chen, Y. Zhang, T. Yu, R. Yao, J. Chen, Q. Fu, J. Zhou, J. Wang, NEIL3 contributes to the Fanconi anemia/BRCA pathway by promoting the downstream double-strand break repair step. Cell Rep 41, 111600 (2022).

50. X. Li, W.-D. Heyer, Homologous recombination in DNA repair and DNA damage tolerance. Cell Res 18, 99–113 (2008).

51. P. Selemenakis, N. Sharma, M. E. Uhrig, J. Katz, Y. Kwon, P. Sung, C. Wiese, RAD51AP1 and RAD54L Can Underpin Two Distinct RAD51-Dependent Routes of DNA Damage Repair via Homologous Recombination. Front Cell Dev Biol 10 (2022), doi:10.3389/fcell.2022.866601.

52. E. Bolderson, N. Tomimatsu, D. J. Richard, D. Boucher, R. Kumar, T. K. Pandita, S. Burma, K. K. Khanna, Phosphorylation of Exo1 modulates homologous recombination repair of DNA double-strand breaks. Nucleic Acids Res 38, 1821–1831 (2010).

53. S. Zona, L. Bella, M. J. Burton, G. Nestal de Moraes, E. W. F. Lam, FOXM1: An emerging master regulator of DNA damage response and genotoxic agent resistance. Biochim Biophys Acta 1839, 1316 (2014).

54. Y. Xie, Y.-K. Liu, Z.-P. Guo, H. Guan, X.-D. Liu, D.-F. Xie, Y.-G. Jiang, T. Ma, P.-K. Zhou, RBX1 prompts degradation of EXO1 to limit the homologous recombination pathway of DNA double-strand break repair in G1 phase. Cell Death Differ 27, 1383–1397 (2020).

55. E. K. Holl, B. P. O’Connor, T. M. Holl, K. E. Roney, A. G. Zimmermann, S. Jha, G. Kelsoe, J. P.-Y. Ting, Plexin-D1 Is a Novel Regulator of Germinal Centers and Humoral Immune Responses. The Journal of Immunology 186, 5603–5611 (2011).

56. R. B. Serafim, C. Cardoso, C. B. Storti, P. da Silva, H. Qi, R. Parasuram, G. Navegante, J. P. S. Peron, W. A. Silva, E. M. Espreafico, M. L. Paçó-Larson, B. D. Price, V. Valente, HJURP is recruited to double-strand break sites and facilitates DNA repair by promoting chromatin reorganization. Oncogene 2024 43:*11* 43, 804–820 (2024).

57. M. Rakhmanov, H. Sic, A. K. Kienzler, B. Fischer, M. Rizzi, M. Seidl, K. Melkaoui, S. Unger, L. Moehle, N. E. Schmit, S. D. Deshmukh, C. K. Ayata, W. Schuh, Z. Zhang, F. L. Cosset, E. Verhoeyen, H. H. Peter, R. E. Voll, U. Salzer, H. Eibel, K. Warnatz, High Levels of SOX5 Decrease Proliferative Capacity of Human B Cells, but Permit Plasmablast Differentiation. PLoS One 9, e100328 (2014).

58. L. Saul, K. M. Ilieva, H. J. Bax, P. Karagiannis, I. Correa, I. Rodriguez-Hernandez, D. H. Josephs, I. Tosi, I. U. Egbuniwe, S. Lombardi, S. Crescioli, C. Hobbs, F. Villanova, A. Cheung, J. L. C. Geh, C. Healy, M. Harries, V. Sanz-Moreno, D. J. Fear, J. F. Spicer, K. E. Lacy, F. O. Nestle, S. N. Karagiannis, IgG subclass switching and clonal expansion in cutaneous melanoma and normal skin. Sci Rep 6, 29736 (2016).

59. M. D. Iglesia, J. S. Parker, K. A. Hoadley, J. S. Serody, C. M. Perou, B. G. Vincent, Genomic Analysis of Immune Cell Infiltrates Across 11 Tumor Types. J Natl Cancer Inst 108, djw144 (2016).

60. J. Li, J. Zheng, R. Zhang, W. Zhang, J. Zhang, Y. Zhang, Pan-cancer analysis based on epigenetic modification explains the value of HJURP in the tumor microenvironment. Scientific Reports 2022 12:*1* 12, 1–10 (2022).

61. G. Tlaposo, H. W. Nijman, ~willem Stoorvogel, R. Leijendekker, C. Hardingfl Cornelis, J. M. Melief, H. J. Geuze, B Lymphocytes Secrete Antigen-presentingVesicles. (available at http://rupress.org/jem/article-pdf/183/3/1161/1678390/1161.pdf).

62. H.-D. Phan, M. N. Longjohn, D. J. B. Gormley, R. H. Smith, M. Dang-Lawson, L. Matsuuchi, M. R. Gold, S. L. Christian, CD24 and IgM Stimulation of B Cells Triggers Transfer of Functional B Cell Receptor to B Cell Recipients Via Extracellular Vesicles. The Journal of Immunology 207, 3004–3015 (2021).

63. C. Rival, M. Mandal, K. Cramton, H. Qiao, M. Arish, J. Sun, J. V. McCann, A. C. Dudley, M. D. Solga, U. Erdbrügger, L. D. Erickson, B cells secrete functional antigen-specific IgG antibodies on extracellular vesicles. Scientific Reports 2024 14:*1* 14, 1–16 (2024).

64. M. F. Gutknecht, N. E. Holodick, T. L. Rothstein, B cell extracellular vesicles contain monomeric IgM that binds antigen and enters target cells. iScience 26, 107526 (2023).

65. G. Restivo, A. Tastanova, Z. Balázs, F. Panebianco, M. Diepenbruck, C. Ercan, B. T. Preca, J. Hafner, W. P. Weber, C. Kurzeder, M. Vetter, S. M. Soysal, C. Beisel, M. Bentires-Alj, S. Piscuoglio, M. Krauthammer, M. P. Levesque, Live slow-frozen human tumor tissues viable for 2D, 3D, ex vivo cultures and single-cell RNAseq. Commun Biol 5 (2022), doi:10.1038/S42003-022-04025-0.

66. L. Chang, G. Zhou, O. Soufan, J. Xia, miRNet 2.0: network-based visual analytics for miRNA functional analysis and systems biology. Nucleic Acids Res 48, W244–W251 (2020).

68. A. P. Liu, J. Ewald, Z. Pang, E. Legrand, Y. S. Jeon, J. Sangiovanni, O. Hacariz, G. Zhou, J. Head, N. Basu, J. Xia, ExpressAnalyst: A unified platform for RNA-sequencing analysis in non-model species. Nat Commun 14, 2995 (2023).

67. T. Li, J. Fu, Z. Zeng, D. Cohen, J. Li, Q. Chen, B. Li, X. S. Liu, TIMER2.0 for analysis of tumor-infiltrating immune cells. Nucleic Acids Res 48, W509–W514 (2020).

68. G. Zhou, O. Soufan, J. Ewald, R. E. W. Hancock, N. Basu, J. Xia, NetworkAnalyst 3.0: a visual analytics platform for comprehensive gene expression profiling and meta-analysis. Nucleic Acids Res 47, W234–W241 (2019).

69. T. Metsalu, J. Vilo, ClustVis: a web tool for visualizing clustering of multivariate data using Principal Component Analysis and heatmap. Nucleic Acids Res 43, W566–W570 (2015).

70. H. Heberle, G. V. Meirelles, F. R. da Silva, G. P. Telles, R. Minghim, InteractiVenn: a web-based tool for the analysis of sets through Venn diagrams. BMC Bioinformatics 16, 169 (2015).

71. Z. Tang, C. Li, B. Kang, G. Gao, C. Li, Z. Zhang, GEPIA: a web server for cancer and normal gene expression profiling and interactive analyses. Nucleic Acids Res 45, W98–W102 (2017).

