## Supplementary data for "B-Cells Extracellular Vesicles Shape Melanoma Response to Immune Checkpoint Therapy"

### Supplementary Figures

#### Fig. S1

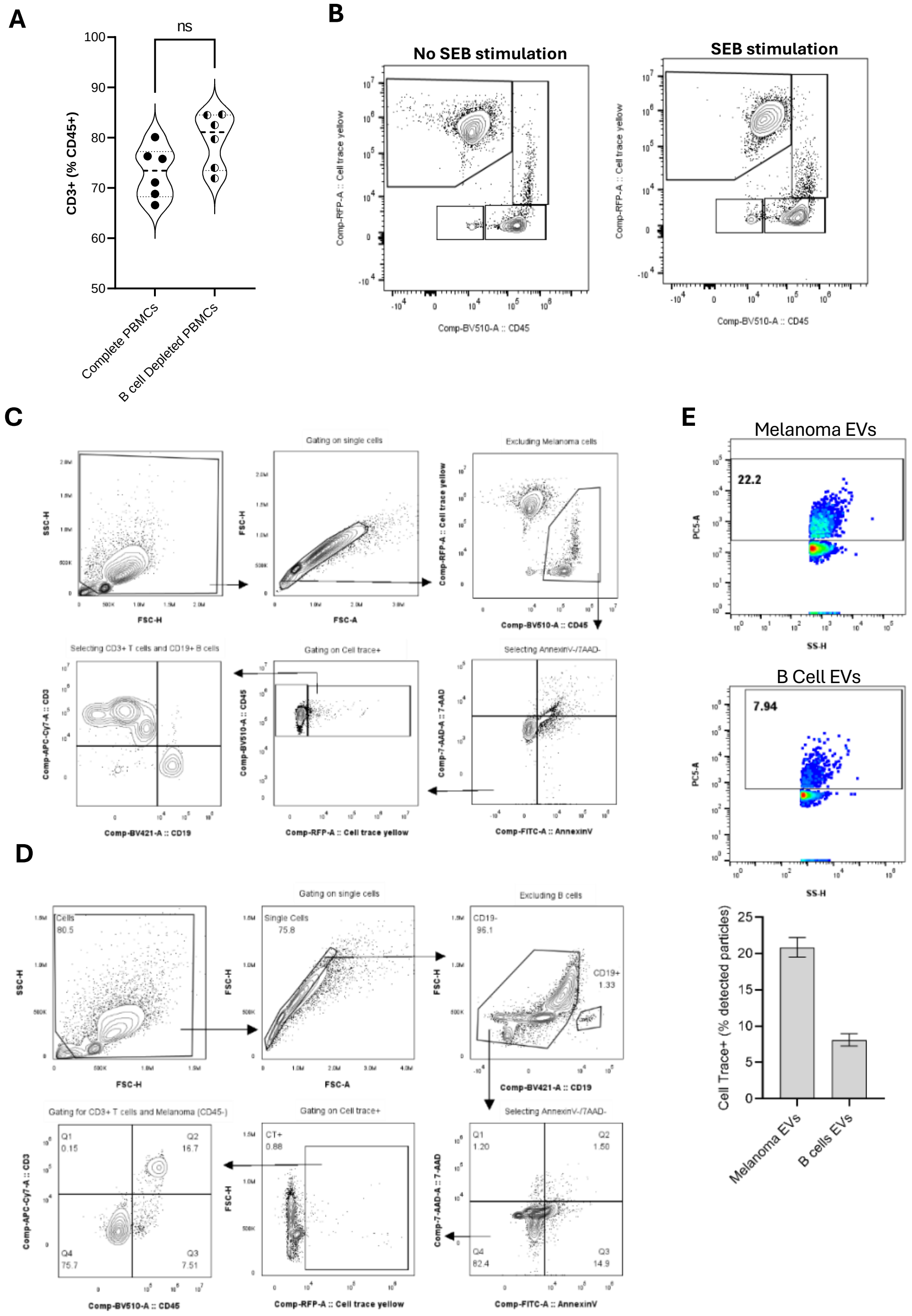

**Figure S1.** (A) $CD3^{+}$ T cells in complete and B cell-depleted cultures, with or without SEB stimulation. Presented as percentage of $CD45^{+}$ population. *P* values were determined by paired t-test (B) Representative FACS plots of cell trace-stained Melanoma co-culture and (C) gating strategy. (D) gating strategy for cell trace-stained B cells co-culture. (E) Representative FACS plot of EVs isolated from cell trace-stained Melanoma cells and cell-trace stained B cells cells, with their quantification. Samples were acquired with nano-analyzer NanoFCM.

**Fig. S2**

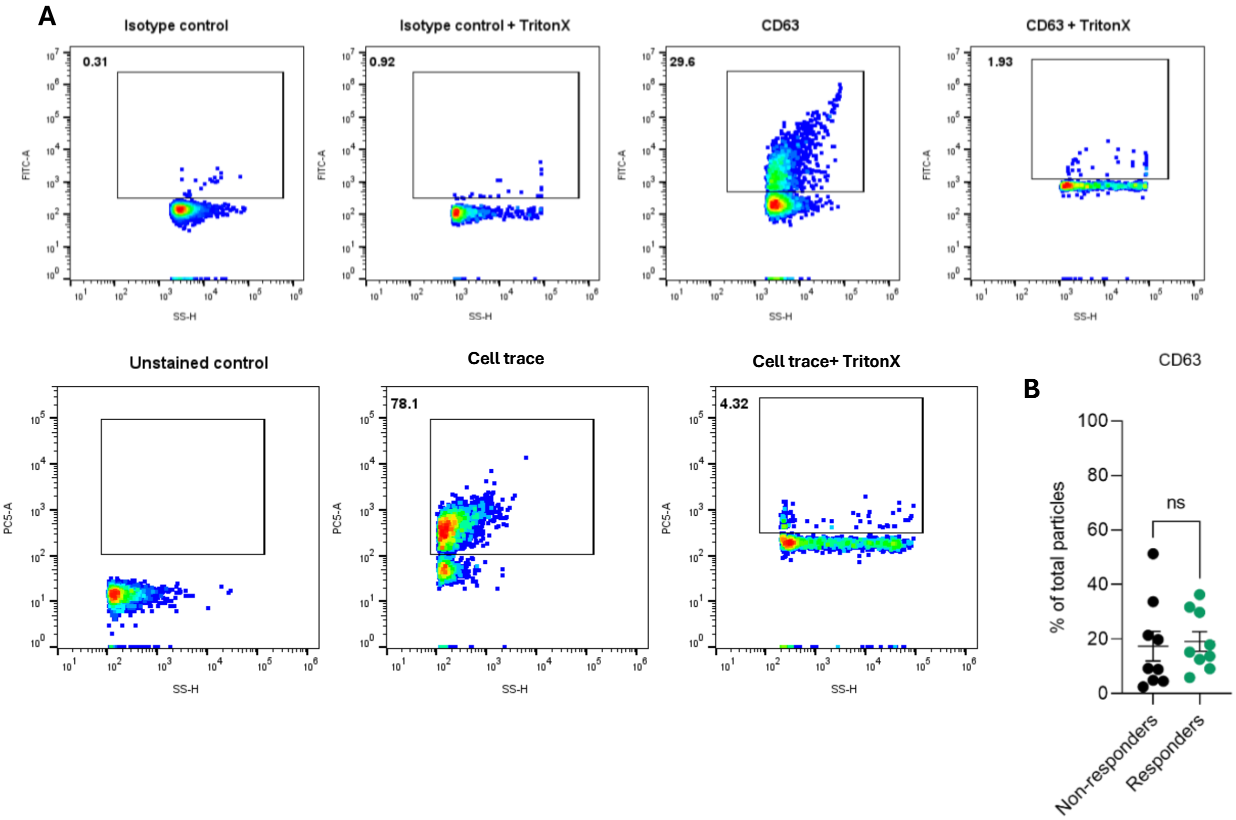


**Figure S2.** (A) Representative FACS plots for EVs detection controls. Unstained control, cell trace to stain EVs membranes, Isotype control, CD63 antibody staining and their respective triton-X controls. Samples were acquired with nano-analyzer NanoFCM. (B) CD63$^{+}$ EVs, presented as percentage of detected particles in responders (green, n=9) and non-responders (black, n=6). P values were determined by Welch's t-test.

**Fig. S3**

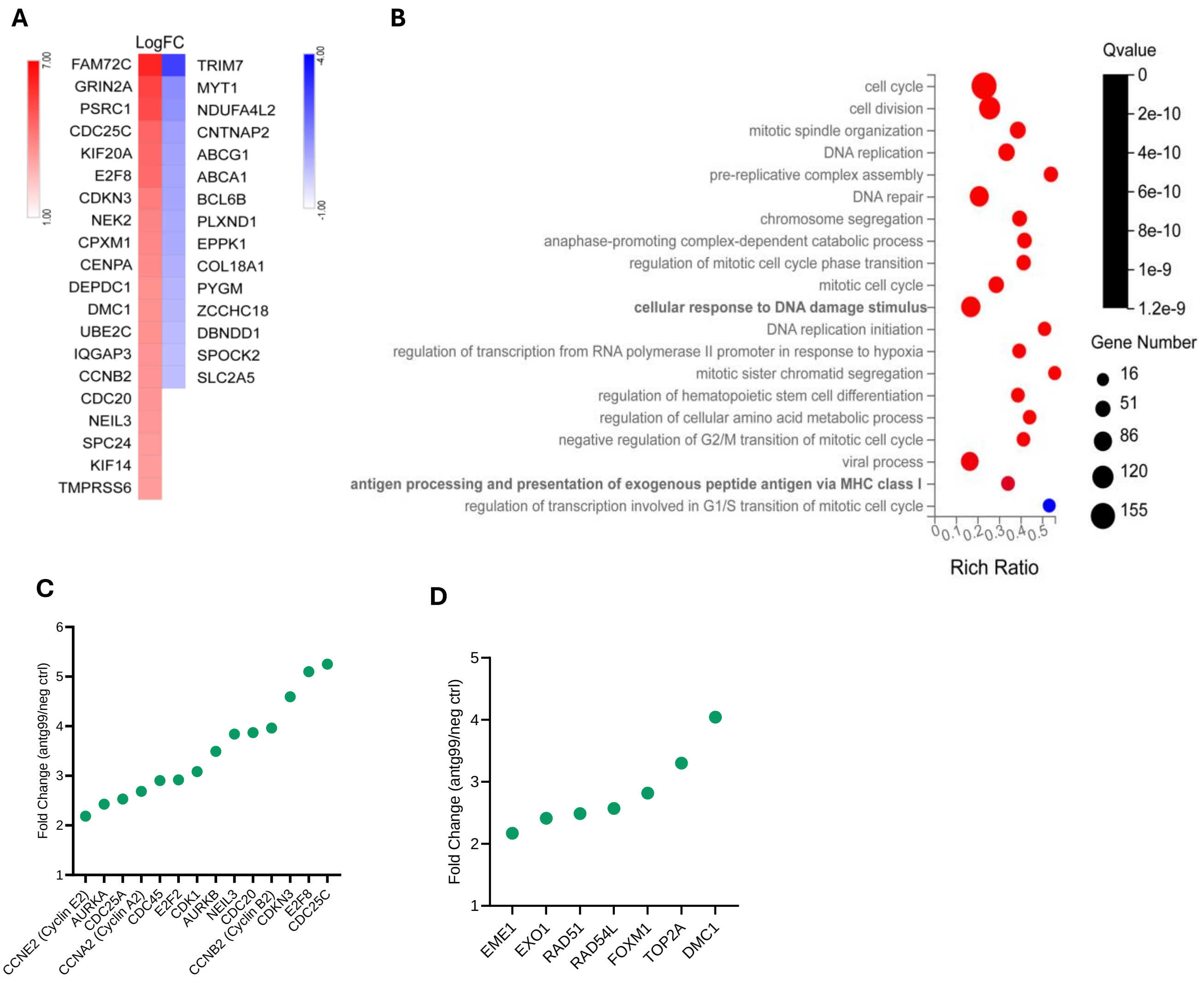


**Figure S3.** (A) Expression heatmap of the top 20 DEGs in antagomiR-99a-5p or negative control antagomiR. (B) bubble chart of enriched biological process terms of significant upregulated genes in antagomiR-99a-5p treated B cells in comparison to negative control antagomiR. Analysis and bubble chart graphs were done using ShinyGO 0.77 (80). (C) Expression of cell cycle genes in antagomiR-99a-5p treated B cells presented as fold change over negative control antagomiR, based on RNAseq. (D) Expression of DDR and HR genes in antagomiR-99a-5p treated B cells presented as fold change over negative control antagomiR, based on RNAseq.
